# Activation of the type 3 secretion system of enteropathogenic *E. coli* leads to remodeling of its membrane composition and function

**DOI:** 10.1101/2020.12.17.423234

**Authors:** Anish Zacharia, Ritesh Ranjan Pal, Naama Katsowich, Chanchal Thomas Mannully, Aida ibrahim, Sivan Alfandary, Raphael Serruya, Amit K Baidya, Sigal Ben-Yehuda, Ilan Rosenshine, Arieh Moussaieff

## Abstract

The cell envelope of gram-negative bacteria is a complex structure, essential for bacterial survival and for resistance to many antibiotics. Channels that cross the bacterial envelope and the host cell membrane form secretion systems that are activated upon attachment to host, enabling bacteria to inject effector molecules into the host cell, required for bacterial-host interaction. The type III secretion system (T3SS) is critical for the virulence of several pathogenic bacteria, including enteropathogenic *E. coli* (EPEC). The EPEC T3SS activation is associated with repression of carbon storage regulator (CsrA), resulting in gene expression remodeling, which is known to affect EPEC central carbon metabolism and contributes to the adaptation to a cell-adherent lifestyle in a poorly understood manner. We reasoned that the changes in bacterial envelope upon attachment to host and the activation of a secretion system may involve a modification of the lipid composition of bacterial envelope. Accordingly, we performed a lipidomics analysis on mutant strains that simulate T3SS activation. We saw a shift in glycerophospholipid metabolism towards the formation of lysophospholipids, attributed to corresponding upregulation of the phospholipase *pldA* and the acyltransferase *ygiH* upon T3SS activation in EPEC. We also detected a shift from menaquinones and ubiquinones to undecaprenyl lipids, concomitant to abnormal synthesis of O-antigen. The remodeling of lipid metabolism is mediated by CsrA and associated with increased bacteria cell size and Zeta potential, and a corresponding alteration in EPEC permeability to vancomycin, increasing the sensitivity of T3SS-activated strains and of adherent wild type EPEC to the antibiotic.

**Importance:** The characterization of EPEC membrane lipid metabolism upon attachment to host is an important step towards a better understanding the shift of EPEC, a notable human pathogen, from a planktonic to adherent life style. It may also apply to other pathogenic bacteria that use this secretion system. We predict that upon attachment to host cells the lipid remodeling upon T3SS activation contributes to bacterial fitness and promotes host colonization, and show that it is associated with increased cell permeability and higher sensitivity to vancomycin.

To the best of our knowledge, this is the first demonstration of a bacterial lipid remodeling due to activation of a secretion system.

## Introduction

The cell envelope of gram-negative bacteria is a complex structure, consisting of a bilayered plasma membrane, a periplasm, and an outer membrane with proteins and lipids (1). This complex is essential for bacterial survival in harsh environments and for resistance to many antibiotics (2). Remarkably, channels that cross the bacterial envelope and the host cell membrane are regulated upon attachment to host, enabling bacteria to inject effector proteins into the host cell, critical for the bacterial-host interaction (3).

Enteropathogenic *E. coli* (EPEC) is a common cause of pediatric diarrhea (4). Upon attachment to the host intestinal epithelium, this pathogen employs a type three secretion system (T3SS) to inject effector proteins into the host cells. The T3SS is encoded within a pathogenicity island termed the locus of enterocyte effacement (LEE) (5), composed of a cluster of transcriptional units containing 41 genes encoding for T3SS structural components, six translocated effectors, and related proteins (6, 7). The LEE5 operon contains three genes: *tir, cesT*, and *eae*, encoding Tir (Translocated intimin receptor), CesT, and intimin, respectively (8, 9). Tir is the most abundant effector and the first to be translocated into the host cell (10, 11). CesT is a homodimer chaperone associated with many effectors, of which Tir is the major one (12, 13). CesT binds to two regions in Tir at the N- and C-terminus through a specific recognition motif, promoting Tir stability and its translocation to the host (12). Intimin, the third product of the LEE5 operon, is an outer membrane protein that promotes adherence of EPEC to the host via interaction with the surface exposed loop of translocated Tir (14, 15). This attachment type was termed intimate attachment (14).

CesT-Tir interaction is essential for Tir translocation into the host and the subsequent intimate attachment action. The second function of CesT-Tir interaction is related to the rearrangement of gene expression upon EPEC-host contact and the consequent Tir translocation (16). In planktonic EPEC, CesT remains bound to Tir and other effectors and the T3SS is not active but fully assembled, ready to immediately translocate the effectors into the host (17).

The secretory activity of T3SS is activated only upon attachment to the host cell and insertion of the translocon (i.e. the EspBD channel) to the host cell membrane. Three proteins, termed “switch proteins”, form a complex that is responsible to prevent effector secretion in non-attached EPEC and promote secretion upon translocon insertion. How this switch complex work is not fully understood. SepD is one of the switch proteins, and in its absence, the switch complex can no longer prevent secretion via the T3SS into the medium regardless of the host cell. Thus, the sepD mutant but not the wild type mimic the T3SS activation and consequence effector secretion leading to liberation of CesT and thus higher level of free CesT. (7, 11, 12, 17-20). The delivery of these effectors into the host liberates CesT, resulting in increased levels of free CesT in the EPEC cytoplasm. The liberated CesT then interacts with an alternative binding partner, the carbon storage regulator A (CsrA) (12, 16, 21). CsrA is an RNA-binding protein and posttranscriptional regulator, which coordinates numerous bacterial functions, including motility, metabolism, and virulence (22, 23). Notably, the elevated levels of free CesT, upon effectors injection, competitively inhibit CsrA-mRNA interaction (12, 16, 21, 24). Since CsrA binds to the mRNA of numerous genes and regulates the stability and/or translation of these mRNAs, CesT-CsrA interaction result in remodeling of gene expression (**Fig. 1**).

**Fig 1.**
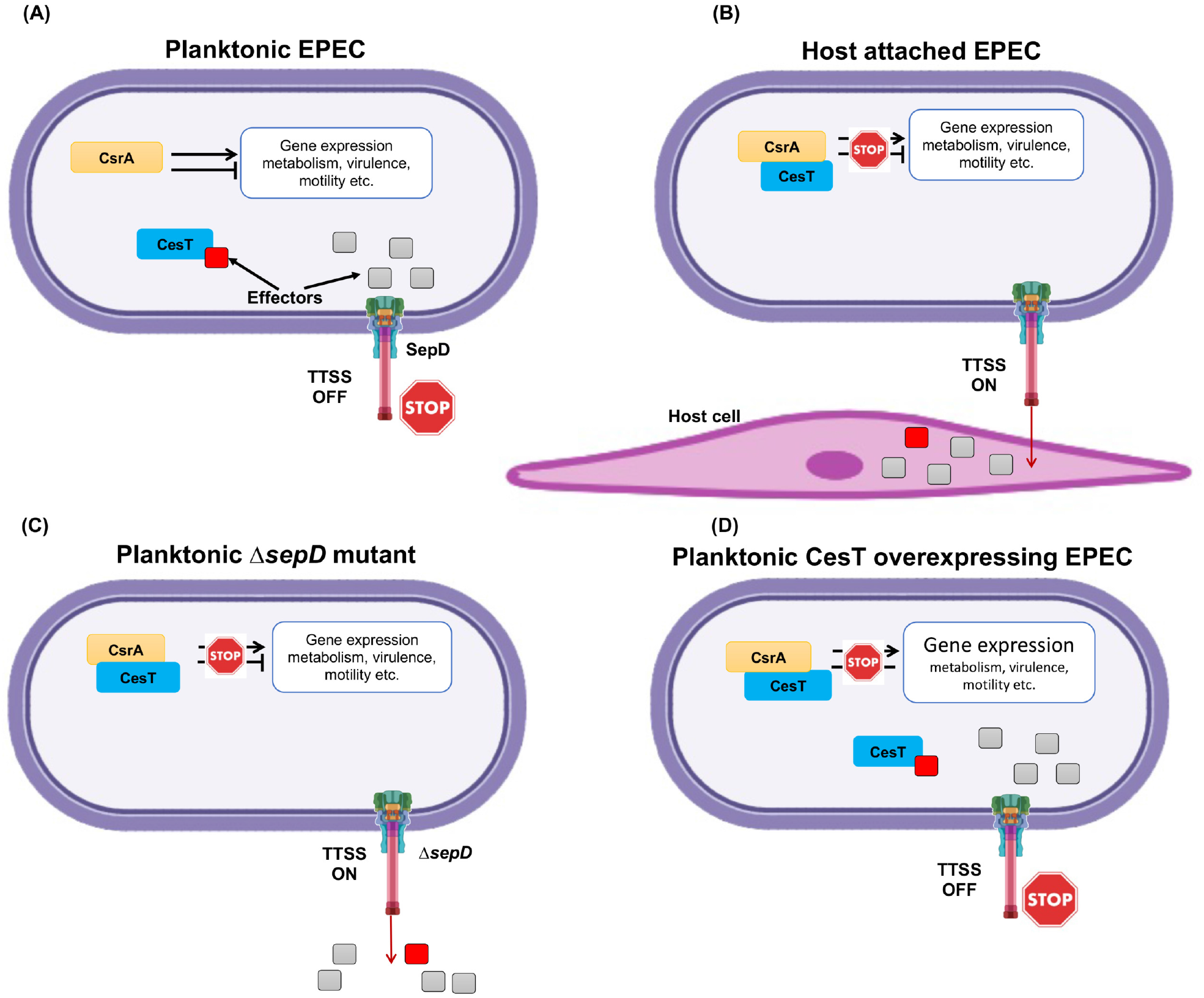
A scheme of the T3SS regulation of EPEC attachment to the host cell. To mimic the attached state of EPEC and study concomitant metabolic shifts we used T3SS regulated strains as following. (A) Planktonic wild type EPEC: The T3SS is not active. The effectors accumulate in the cytoplasm, and some of them (indicated as a red square) bind to CesT and sequester it. In these bacteria, CsrA is free to bind mRNA and regulate gene expression. (B) Host-attached wild type EPEC: The T3SS is activated upon contact with the host, eliminating the effectors from the EPEC cytoplasm by injecting them into the host cell. The liberated CesT then binds to CsrA to inhibit CsrA-mRNA interaction. (C) Planktonic Δ*sepD* mutant: The T3SS is constitutively active, and effectors are secreted. The liberated CesT binds to CsrA and inhibits CsrA-mRNA interaction. (D) Planktonic CesT-overexpressing EPEC: CesT is expressed at levels that allow binding to all the CesT-binding effectors and CsrA, resulting in inhibition of CsrA-mRNA interactions regardless of T3SS activity.

Importantly, while wild type and sepD mutant have similar total CesT levels, the levels of free CesT varies. Consequently, the levels of CsrA inhibition by CesT also varies between the two strains (16, 25). As to the CesT overexpressing strain– this strain has higher levels of total CesT and thus, higher level of free CesT regardless of host cell or secretion of effectors.

The significance of the CesT-CsrA switch to the bacterial physiology can be partially extrapolated from studies comparing wild type *E. coli* to *csrA* mutants (22, 26-29). However, *csrA* mutants are not an ideal model to mimic host interacting EPEC since the CesT-CsrA interaction induces only partial and temporal CsrA inhibition. Previous reports point to alternative approaches that mimic the physiological state of host-attached EPEC. The first approach employs the EPEC *sepD* mutant (Δ*sepD*), which expresses T3SS that constitutively secretes Tir regardless of host attachment (16, 30). Thus, the *sepD* mutant contains higher levels of liberated CesT, that is free to interact with CsrA. Another approach uses wild-type EPEC containing a plasmid that expresses CesT under an isopropyl-D-thiogalactopyranoside (IPTG) regulated promoter (WT/pCesT). Upon IPTG treatment it overexpresses CesT that readily interacts with CsrA (**Fig. 1**).

Here, we combined lipidomics and genetic methodologies to characterize the shift of EPEC metabolism under conditions that mimic infection and the associated CsrA inhibition. We sought to unveil possible shifts in membrane lipid metabolism that take place upon the secretion system activation, by an unbiased lipidomics analysis. However, obtaining biological material from injecting EPEC in quantities that allow lipidomics analysis is challenging, as only a small subpopulation is engaged in injection, while in the rest of the population that are not in direct contact with the host, the T3SS remains inactive (11). To overcome this hurdle and obtain uniform populations, we compared wild type EPEC that does not secrete Tir in a planktonic state (thus Tir sequesters CesT), to an isogenic EPEC Δ*sepD* mutant that constitutively secretes Tir, and consequently, CesT is liberated, free to interact with CsrA.

Our lipidomics analysis pointed to a major metabolic signature of the shift from planktonic to adherent life style (**Fig. S1**). We expect this lipid shift to be a general trait for secretion systems, in particular in *E. coli* spp. experiencing CsrA inhibition. We assume that this lipid modulation is involved in the adaptation of EPEC to the cell-adherent lifestyle.

## Results

We used EPEC strains that mimic the host-attached status to study the changes in their lipid metabolism. We compared wild type EPEC that does not secrete Tir in a planktonic state (thus Tir sequesters CesT), to an isogenic EPEC Δ*sepD* mutant that constitutively secretes Tir, and consequently, CesT is liberated, free to interact with CsrA. To verify the CsrA-mediated regulation of metabolism in these strains, we used EPEC with a deleted *csrA* gene (Δ*csrA*). The latter was also used as a reference to available data of the metabolic profile of the EPEC and *E. coli* K12 Δ*csrA* mutant (28, 31, 32). We grew the bacteria under conditions optimal for the expression of the T3SS genes (i.e., growth in 500 mL DMEM, 37°C, without shaking to optical density (OD)_600_ 0.6). Under these experimental conditions, Δ*csrA* strains showed an attenuated growth rate (**Fig. S2)**, in line with previous literature (23, 28, 32). The Δ*sepD*Δ*cesT* double mutant showed a mild attenuation in growth rate, whereas the Δ*sepD* strain and EPEC supplemented with CesT expressing plasmid (WT/pCesT) showed growth rates similar to that of the wild type.

### CesT-CsrA interaction confers alterations in PL and terpenoid-quinone pathways

To study potential changes in EPEC lipid composition following T3SS activation, we performed a lipidomics analysis. We focused on the Δ *sepD* strain, and used the WT/pCesT, overexpressing CesT, to confirm that the changes seen in the Δ *sepD* strain are CesT-dependent. A Δ *sepD*,Δ *cesT*, double mutant, assumed to suppress the Δ *sepD* mutation, was also used for further confirmation.

A total of 15,827 metabolic features were detected. After the exclusion of possible artifact features (see Methods section), downstream lipidomics analysis was carried out using 3,277 metabolic features. We first analyzed the lipid composition of wild type vs the *sepD* mutant. Principal component analysis (PCA) demonstrated a striking separation between the lipidome of the Δ *sepD* mutant and that of wild type EPEC, with PC1 accounting for 97.9% of the variance (**Fig. 2A**), suggesting that the increased levels of free CesT in the Δ *sepD* mutant induced a considerable shift in lipid metabolism. To test this hypothesis, we compared the lipidome of wild type EPEC and *sepD* mutant to that of EPEC double mutant Δ *sepD*,Δ *cesT*. The PCA of the lipidome of EPEC Δ *sepD*Δ *cesT* showed no shift in the lipid composition compared to the wild type strain (**Fig. 2B**), indicating that CesT is required for inducing the shift in the lipid composition. To reinforce this premise we tested wild type EPEC overexpressing CesT (WT/pCesT). In this case, the overexpression of CesT was sufficient for inducing a shift in lipid composition. This shift was more pronounced than that in the Δ *sepD* mutant (**Fig. 2B**, principal component 1 of the PCA), likely due to higher levels of free CesT in the overexpressing bacteria compared to these in the Δ *sepD* mutant. Taken together, our data suggest that an increase in the levels of CesT is necessary and sufficient to induce a considerable shift in the lipid composition of EPEC.

**Fig 2.**
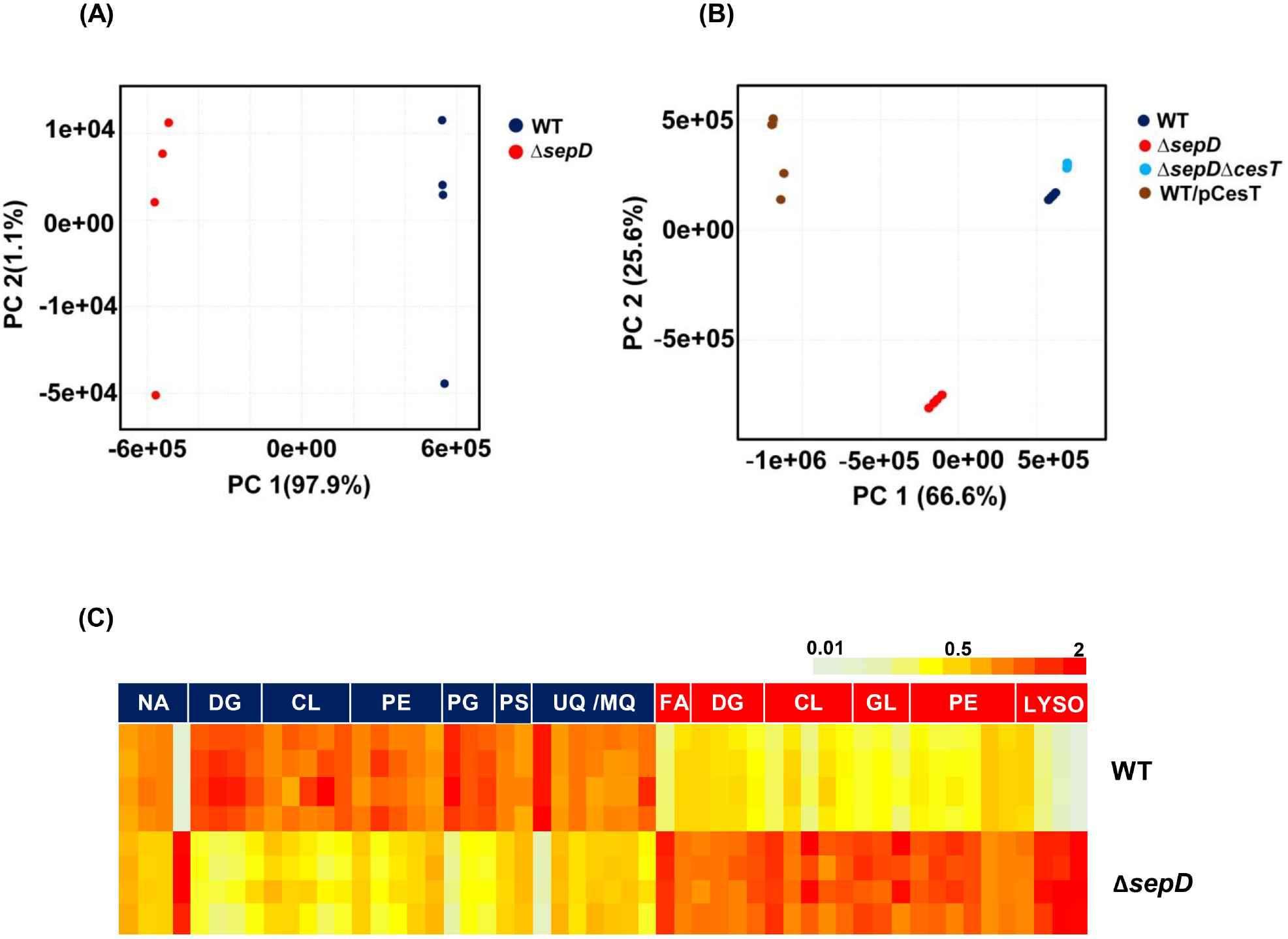
Levels of free CesT correlate with a profound effect on EPEC lipid metabolism. (A) Principal component analysis (PCA) of lipids in wild type (WT) vs. ΔsepD EPEC (N=4). (B) PCA of the lipidome of WT, ΔsepD, ΔsepDΔcesT and WT/pCesT. (C) A heat map of the differential lipids (Variable Importance in Projection (VIP) >1) in sepD mutants, divided into lipid subclasses. Lipid subclasses with upregulated accumulation in mutants are in red and lipid subclasses with downregulated accumulation in mutants are in blue quadrants. Color in each column indicates the fold change from mean abundance of the particular feature across all samples. Lyso, lysophospholipids; GL, Glycolipids; CL, Cardiolipins; DG, Diacylglycerols; FA, Fatty acids; Q/MK, Ubiquinone/Menaquinone; PE, phosphatidylethanolamines; PG, phosphatidylglycerols; PS, phosphatidylserines; NA, not assigned.

A Variable Importance in Projection (VIP) analysis of Δ *sepD* vs wild type EPEC lipidome revealed 54 differential (VIP>1) mass features. Phospholipids (PLs) and isoprenoids of the quinone terpenoid subclass were predominantly discriminant features in this analysis (**Fig. 2C**). Phosphatidylethanolamine (PE) and phosphatidylglycerol (PG) are the major lipid components of bacterial membranes. The inner membrane of Escherichia coli contains 70–80% PE and 20–25% PG (see Figure S1 for structures). Within the differential lipids, all 4 lysophospholipids (LysoPLs; phospholipids with a single carbon chain and a polar head group) were highly accumulated in the *SepD* mutants. The distribution of PLs in the differential lipids was more complex. All 6 upregulated PLs were PEs, while out of the 10 PLs in the group of lipids that were downregulated in *SepD* mutants 5 were PEs, 3 PGs and 2 phosphatidylserine (PS). All menaquinone and ubiquinone species were downregulated in the *SepD* mutants. Given these results, we focused our analyses thereof on the PL and terpenoid-quinones pathways, and identified lipids of these two main differential classes from the whole dataset (**Table S1**).

### T3SS activation drives a remodeling of the glycerophospholipid pathway

Our VIP analysis pointed to PLs and quinone terpenoids as major differential lipids classes following T3SS activation, and implied to a metabolic shift from PGs in the glycerophospholipid pathway. To better define the shifts in the pathway, we quantified the abundance of all identified phospholipid species per subclasses, using our lipidomics analysis data. We identified 41 PE species and 12 PGs. We observed a shift in the PL composition. This shift was further verified using a second dataset, where each PL subclass was quantified with a corresponding heavy isotope internal standard. All data presented herein is of the latter quantified dataset. We found a considerable decrease in the abundance of PGs in Δ*sepD* and WT/pCesT EPEC (**Fig. 3A**), whereas mild increase in that of PEs in WT/pCesT EPEC (**Fig. 3B**). Aligned with our VIP analysis, we also revealed a higher abundance of the identified LysoPLs in Δ*sepD* and WT/pCesT EPEC (34 identified lipid species; (**Fig. 3C**)). Cardiolipins (CLs; 10 identified lipid species; (**Fig. 3D**)) were upregulated in the mutant strain as well. To pinpoint the enzymes involved in the shift in PL metabolism, and in particular the dramatic increase in the abundance of lysoPLs in the Δ*sepD* and WT/pCesT strains, we performed a network-based analysis of Δ*sepD* PL metabolism (**Fig. 3E**). This analysis underscored two enzymes: PldA that encodes for phospholipase, and YgiH, that functions as a glycerol-3-phosphate acyltransferase for lysoPL biosynthesis (33). We then quantified the expression of the corresponding genes by q-RT-PCR, and noted a considerable upregulation in the RNA levels of these genes upon T3SS activation (**Fig. 3F-G**). These results are in alignment with the increased concentrations of the corresponding lipids.

**Fig 3.**
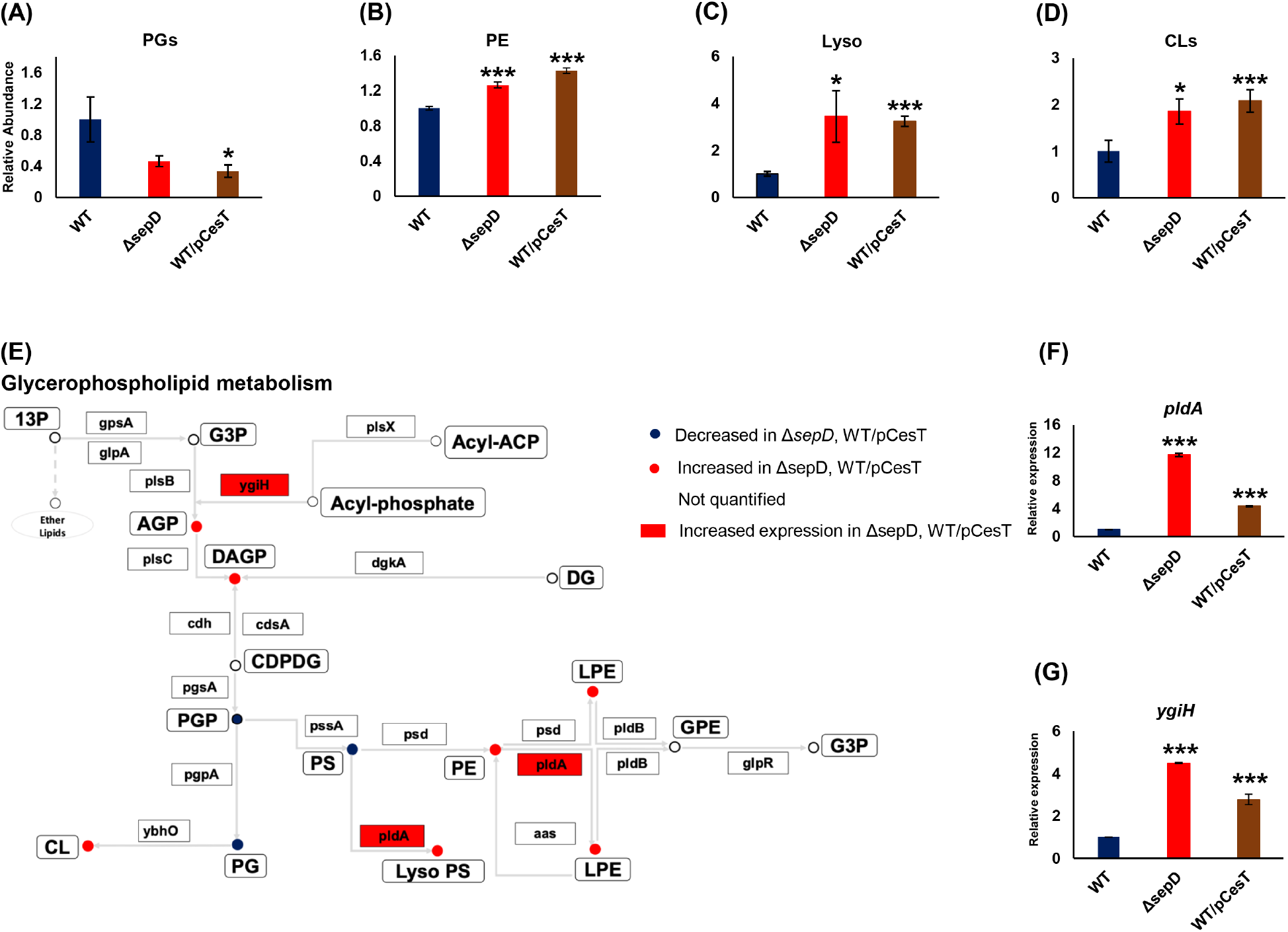
Repression of CsrA by CesT leads to a shift from phosphatidylglycerols (PGs) to phosphatidylethanolamines (PEs) and lysophospholipids (LysoPLs) on the one hand and CLs on the other. (A) The relative total abundance of identified PGs (12 identified lipid species; A); PEs (41 identified lipid species; B); LysoPLs (34 identified lipid species; C), and cardiolipins (10 identified lipid species; CLs; D) is presented. (E) A metabolic map of phospholipids was adapted from the Kyoto Encyclopedia of Genes and Genomes (KEGG) for Escherichia coli O127:H6 E2348/69 (EPEC), and modified. Lipid classes are represented by circles, with their names in white quadrants. Lipids abbreviations: 13P, Glycerone phosphate; G3P, Glycerol-3-phosphate; AGP, Acylglycerol-3-phosphate; DAGP, 1,2-Diacylglycerol 3-phosphate; CDPDG 1,2-Diacylglycerol-cytidine-5-diphosphate; GPE, Glycerophosphoethanolamine; LPE, Lysophosophoethanolamine; LysoPS, Lysophosphoserine; PGP, Phosphatidylglycerophosphate; PE, Phosphatidylethanolamine; PS, Phosphatidylserine; PG, Phosphatidylglycerol. Enzymes are represented by arrows, with their numbers in quadrants. Enzymes common nomenclature is used in the scheme. For an unambiguous identification, enzyme nomenclature (EC number system) is hereby given, along with further commonly used names: [1.1.1.94]-gpsA, glycerol-3-phosphate dehydrogenase; [1.1.5.3]- glpD, sn-glycerol-3-phosphate dehydrogenase; [2.3.1.15]- plsB, glycerol-3-phosphate O-acyltransferase; [2.3.1.274]- plsX, phosphate acyltransferase; [2.3.1.275]- plsY (ygiH), glycerol-3-phosphate acyltransferase; [2.3.1.51]- plsC (parF), 1-acyl-sn-glycerol-3phosphate acyltransferase; [2.7.1.107]- dgkA, diacylglycerol kinase; [3.6.1.26]- cdh, CDP-diacylglycerol phosphotidylhydrolase; [2.7.7.41]- cdsA, (cds) CDP-diglyceride synthase; [EC:2.7.8.5]- pgsA, phosphatidylglycerophosphate synthetase; [3.1.3.27]- pgpA (yajN), phosphatidylglycerophosphatase A; [2.7.8.8]- pssA (pss), phosphatidylserine synthase; [3.1.1.32]- pldA, outer membrane phospholipase A; [<u>4.1.1.65</u>]- psd, phosphatidylserine decarboxylase; [3.1.1.4]- pldA, outer membrane phospholipase A; [3.1.1.5]- pldB, lysophospholipase L; [3.1.4.46]- glpQ (ugpQ), glycerophosphodiester phosphodiesterase. The relative expression of pldA and ygiH in the EPEC strains were quantified using quantitative PCR (qPCR) (F-G). N=4 for all data, besides the RT-qPCR, where N=3. Data are presented as mean ± SE. *, p<0.05; ***, p<0.001.

Assuming that the changes we identified in PL metabolism may result in reduction of membrane integrity, we stained the membrane by the liphophilic dye, FM4-64, and found a lower uptake of the dye in Δ*sepD* and CesT overexpression strains (**Fig. S3**).

To assess the involvement of CsrA in the T3SS-dependent shift we identified in PL metabolism, we first mined a *csrA*-mutant transcriptome database (24) for the glycerophospholipid pathway genes. We integrated our lipidomics data and the published gene expression (RNA-seq) data for a better understanding of the PL network. The mined gene expression data was well-aligned with the results of our analyses of the abundance of PL subclasses (**Fig. S4A**). To verify that the shift in PL metabolism is mediated by inhibition of CsrA by CesT we quantified the PL subclasses in EPEC Δ*csrA* mutant and compared it to Δ*sepD* and wild type strains. We found in both Δ*csrA and* Δ*sepD* a similar shift from PGs to PEs and increased levels of LysoPLs (**Fig. S4B-D**). We then evaluated the expression of *pldA* and *YgiH* and could detect a substantially higher expression in Δ*csrA and* Δ*sepD* compared to wild type EPEC (**Fig. S4E-G**). These results further suggest that the shift in PL metabolism is mediated via CsrA inhibition by CesT. The growth conditions in our study were set to mimic infection conditions (16, 25). Nevertheless, our gene expression analysis was in agreement with the mined data, although these data was obtained using EPEC grown under different conditions. Altogether, our results point to a CsrA-mediated regulation of PL metabolism in EPEC, and suggest that this modulation is maintained in different growth conditions.

### Undecaprenyl lipid biosynthesis is upregulated, whereas synthesis of menaquinones and ubiquinones is downregulated upon T3SS activation

Isoprenoids (“terpenoids”) are structurally diverse lipids with skeletons composed of 5 carbon units. These units are formed from isopentenyl diphosphate (IPP) and dimethylallyl diphosphate (DMAPP) precursors, and are successively assembled (34) (**Figure S1**). All isoprenoids are synthesized by sequential head-to-tail 1′-4 condensations. Undecaprenyl phosphate is an essential long-chain isoprene lipid involved in the biogenesis of bacterial cell envelope carbohydrate polymers including peptidoglycan and O-antigen. It forms a membrane-bound carrier for the assembly of O-antigen units (35, 36). After synthesis, O-antigen is ligated to the core-lipid A, biosynthesizing LPS (35). IPP is also the source of the prenyl side chains of terpenoid quinones - menaquinone and ubiquinone (37).

Our lipidomics analyses suggested that quinone terpenoids biosynthesis is altered in *sepD* mutants (**Fig. 2C**). We thus used the dataset generated by this lipidomics analysis to compare the levels of identified isoprenoid species, per subclasses. This analysis pointed to increased accumulation of end products of undecaprenyl in Δ*sepD* to those of a wild type strain, while decreased ubiquinone and menaquinone end products accumulation, and suggested a shift towards undecaprenyl biosynthesis in the *sepD* mutants (**Fig. 4A-D**). This shift was recapitulated and was, in fact, more pronounced in EPEC overexpressing CesT (**Fig. 4A-D**).

**Fig 4.**
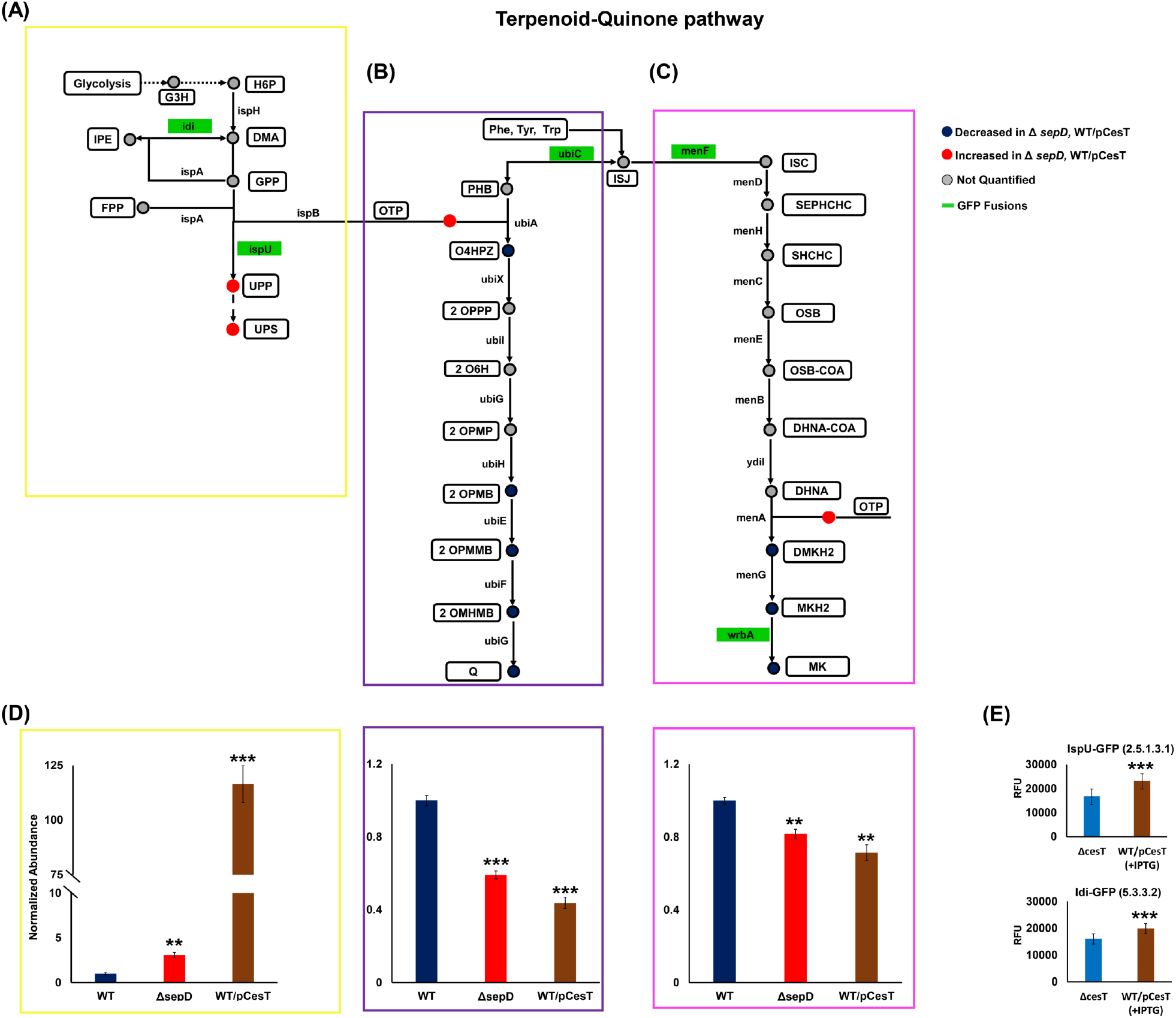
Mimicing the activation of T3SS in EPEC upregulates the biosynthesis of undecaprenyl lipids and downregulates the biosynthesis of menaquinones and ubiquinones. (A-C) Terpenoid quinones pathway was adapted from the Kyoto Encyclopedia of Genes and Genomes (KEGG) for Escherichia coli O127:H6 E2348/69 (EPEC), and modified to include the undecaprenyl, ubiquinone and menaquinone branches of terpenoid pathway, resulting in a simplified scheme of the metabolic network of terpenoid quinone lipids. We divided the map into: (A) undecaprenyl (yellow box), (B) ubiquinone (purple box), and (C) menaquinone (pink box) lipids. The colors of lipid species (in circles) represent their relative accumulation according to our analyses. Lipid species and subclasses are represented in white boxes, and green boxes represent enzyems fused to GFP. Lipids abbreviations: H6P, 1-Hydroxy-2-methyl-2-butenyl- 4- diphosphate; DMA, Dimethylallyl diphosphate; IPE, Isopentenyl diphosphate; GPP, Geranyl diphosphate; FPP, Farnesyl diphosphate; UPP, Undecaprenyl diphosphate; UPS, undecaprenyl species OTP, Octaprenyldiphosphate; PHB, Hydroxybenzoic acid; ISJ, Chorismate; O4HPZ, 4-Hydroxy-3-polyprenyl benzoate; 2OPPP, 2-Octaprenyl phenol; 2O6H, 2-Octaprenyl-6-hydroxyphenol; 2OPMP, 2-Octaprenyl-6-methoxy phenol; 2OPMB, 2-Octaprenyl-6-methoxy- 1,4-benzoquinone; 2OPMMB, 2-Octaprenyl-3-methyl-6-methoxy-1,4-benzoquinone; 2OMHMB, 2-Octaprenyl-3- methyl-5-hydroxy-6-methoxy-1,4-benzoquinone; Q, Ubiquinone; ISC, Isochorismate; SEPHCHC, 2-Succinyl-5- enolpyruvyl-6-hydroxy-3-cyclohexene-1-carboxylate; SHCHC, 6-Hydroxy-2-succinylcyclohexa-2,4-diene-1- carboxylate; OSB, 2-Succinylbenzoate; OSB-COA, 2-Succinylbenzoyl-CoA; DHNA-CoA, 1,4-Dihydroxy-2- naphthoyl-CoA; DHNA, 1,4-Dihydroxy-2-naphthoate; DMKH2, Demethylmenaquinol; MKH2, Menaquinol; MK, Menaquinone. Enzymes are represented by arrows, with their names in quadrants. Enzymes’ common nomenclature is used in the scheme. For an unambiguous identification, enzyme nomenclature (EC number system) is hereby given, along with further commonly used names: [<u>1.17.7.4</u>]- ispH (yaaE, lytB), 4-hydroxy-3-methylbut-2-enyl diphosphate reductase; [2.5.1.1, 2.5.1.10]- ispA, farnesyl diphosphate synthase; [2.5.1.90]- ispB (cel, yhbD), all-trans-octaprenyl- diphosphate synthase; [2.5.1.31]- ispU (uppS, rth, yaeS), ditrans,polycis-undecaprenyl-diphosphate synthase; [4.1.3.40]- ubiC, chorismate lyase; [2.5.1.39]- ubiA, 4-hydroxybenzoate octaprenyltransferase; [2.5.1.129]- ubiX (dedF), 3- octaprenyl-4-hydroxybenzoate carboxy-lyase; [<u>1.14.13.240</u>]- ubiI (visC), 2-methoxy-6-(all-trans-octaprenyl)phenol 4- hydroxylase; [2.1.1.222, 2.1.1.64]- ubiG (pufX, yfaB), bifunctional 3-demethylubiquinone-8 3-O-methyltransferase and 2-octaprenyl-6-hydroxyphenolmethylase; [1.14.13.-]- ubiH (acd, visB) 2-octaprenyl-6-methoxyphenol 4-hydroxylase; [2.1.1.201]- ubiE (yigO), bifunctional 2-octaprenyl-6-methoxy-1,4-benzoquinol methylase and demethylmenaquinone methyltransferase; [1.14.99.60]- ubiF (yleB), 2-octaprenyl-3-methyl-6-methoxy-1,4-benzoquinol oxygenase, 2- octaprenyl-3-methyl-6-methoxy-1,4-benzoquinol hydroxylase; [2.1.1.222, 2.1.1.64]- ubiG (pufX, yfaB) bifunctional 3- demethylubiquinone-8 3-O-methyltransferase and 2-octaprenyl-6-hydroxyphenol methylase; [5.4.4.2]- menF (yfbA), isochorismate synthase 2; [2.2.1.9]- menD, 2-succinyl-5-enolpyruvyl-6-hydroxy-3-cyclohexene-1-carboxylatesynthase; [<u>4.2.99.20</u>]- menH (yfbB), 2-succinyl-6-hydroxy-2,4-cyclohexadiene-1-carboxylate synthase; [4.2.1.113]- menC, o- succinylbenzoate synthase; [6.2.1.26]- menE, o-succinylbenzoateCoA ligase; [4.1.3.36]- menB, 1,4-dihydroxy-2- naphthoyl-CoA synthase; [<u>3.1.2.28</u>]- ydiI (menI), 1,4-dihydroxy-2-naphthoyl-CoA hydrolase; [2.5.1.74]- menA (yiiW), 1,4-dihydroxy-2-naphthoate octaprenyltransferase; [2.1.1.163, 2.1.1.201]- ubiE (menG, yigO), demethylmenaquinone methyltransferase /2-methoxy-6-octaprenyl-1,4-benzoquinol methylase; [1.6.5.2]- wrbA(ytfG), quinone oxidoreductase. (D) Accumulated abundance of undecaprenyl, ubiquinones, and menaquinones lipids, respectively, following accumulation of CesT in the bacteria. (E) GFP flourescence was measured in the strains with null cesT and WT/pCesT. Their expression was determined in terms of relative fluorescence units. Data are presented as mean ± SEM. (N=4). **, p<0.01; ***, p<0.001.

We then tested the influence of CesT levels on the production of key enzymes involved in the terpenoid-quinones biosynthesis. To this end, we fused a GFP reporter gene in frame to the 3’ end of a selected number of genes in their native chromosomal location without perturbing their transcriptional unit. Thus, the GFP levels provide a means to measure enzyme production. We selected for tagging genes whose *gfp*-tagging was previously found to be tolerated by the bacteria (38). Using this approach, we *gfp*-tagged the following genes: *idi*, which forms isopentenyl pyrophosphate, an isoprenoid precursor; *ispU*, involved in the biosynthesis of undecaprenyl pyrophosphate; *ubiC* and *menF*, bridging between menaquinone and ubiquinone pathways, and *wrbA* encoding for menaquinone biosynthesis. To enhance the signal/noise ratio we constructed the GFP reporters in WT/pCesT and Δ*cesT* strains, grown statically in DMEM at 37°C. We detected upregulation of Idi-GFP and IspU-GFP levels upon *CesT* overexpression (**Fig. 4E**). This increase is in agreement with our lipidomics analyses. Changes in the levels of GFP-tagged UbiC, MenF and WrbA, involved in menaquinone and ubiquinone production, were not significant (**Fig. S5**).

Our lipidomics analyses and the quantification of the expression of *gfp*-tagged *idi* and *ispU* are well-aligned with the data we mined on the expression of genes of the quinone terpenoids pathways in *Δ csrA* EPEC (**Fig. S6**).

Taken together, our analyses point to a metabolic shift towards undecaprenyl species upon T3SS activation. Importantly, undecaprenylpyrophoshate (UPP) is required for the synthesis of O-antigen repeating units on surface of the inner leaflet of the inner membrane, and for the flipping of the UPP-O-antigen complex to the outer leaflet, where it is used as a precursor for the biosynthesis of the LPS O-antigen and O-antigen capsule (39). We assumed that the increase in the level of undecaprenyl species we monitored in *SepD* mutants will be reflected in higher biogenesis of O-antigen. To define possible alterations in O-antigen levels, we carried out Western blot analysis of O-Antigen repeats, which suggested increased levels of LPS O-antigen in the Δ*sepD* mutant and *CesT* overexpressing EPEC strains (**Fig. 5A**). Immunofluorescence microscopy demonstrated increased levels of O-antigen (LPS and/or capsular) on the surface of Δ*sepD* mutant and *CesT* overexpressing EPEC strains **(Fig. 5B-C)**, in concordance with the total O-antigen levels observed in Western blot analysis.

**Fig 5.**
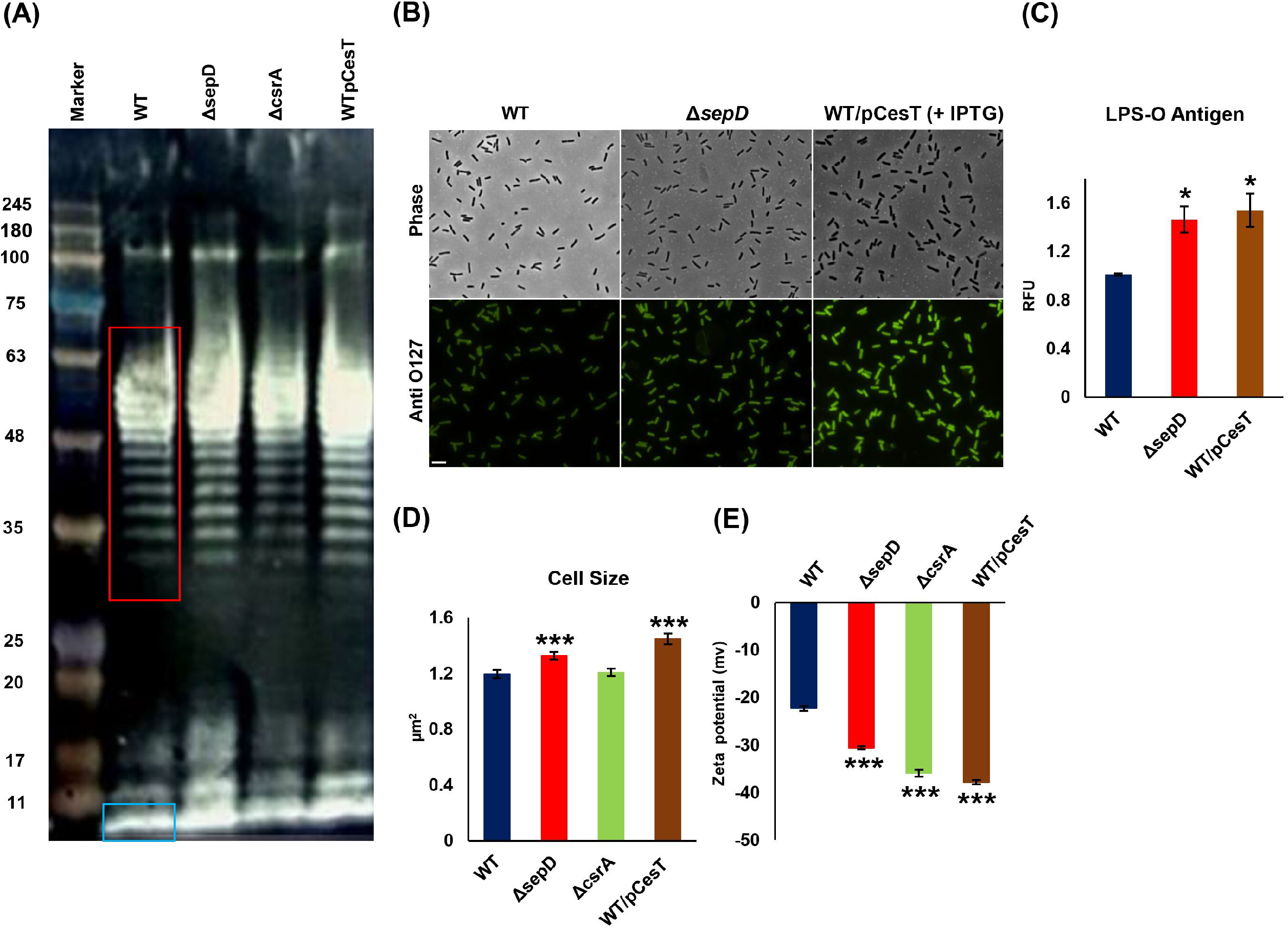
O-Antigen levels, cell size and charge potential in EPEC increase upon T3SS activation. EPEC strains were subcultured in DMEM at 37°C to an OD_600_ of 0.6. (A) LPS was extracted and subjected to Western blotting. O- antigen repeats are boxed in red and the core is in blue box. (B-C) Bacteria were fixed, incubated with anti-O127 antibody, and visualized under phase contrast and fluorescent microscopy. Scale bar indicates 5 μm. The fluorescence intensity was measured using NIS elements AR 4.3 and the relative fluorescence was determined. The data are presented as mean± SEM. (N=3). (D-E) EPEC strains were subcultured in DMEM at 37°C to an OD_600_ of 0.6. Bacteria cell size (N=150; D) was measured under a microscope using Image J software. (E) Zeta potential was measured using Zetasizer Nano ZS. Zeta potential measurements for each sample and the mean charge were recorded (N=12). Data are presented as mean ± SEM. (N=3). *, p<0.05; ***, p<0.001.

We reasoned that the changes we found in the composition of EPEC membrane lipids should be reflected in its physiological and morphological characteristics, and measured the cell size of the Δ*sepD* mutant and WT/pCesT. We found significant increase in their size (**Fig. 5D**), consistent with previous observation made in *csrA* mutant *E coli* (23). Surprisingly, however, the size of the *csrA* mutant was similar to that of wild type EPEC in our hands. The O-antigen content and composition may influence cell surface charge, which plays a role in adhesion of bacteria to surfaces. We saw an increase in the negative charge of the T3SS activated strains (**Fig. 5E**).

The outer and inner membranes form a barrier that separates and protects the bacteria from the extra-cellular environment. Given the profound changes we observed in the lipid composition of the EPEC membranes upon activation of the T3SS, we sought to study the influence of the T3SS on membrane permeability by using a vancomycin permeability-mediated resistance assay (40). We detected higher sensitivity to vancomycin in Δ*sepD* and WT/pCesT EPEC compared to the wild type strain (**Fig. 6A**). To confirm the physiological relevance of the changes in membrane permeability, we wished to evaluate the permeability of adherent vs planktonic wild type EPEC. For this aim, we monitored adherent NleA-GFP EPEC in co-culture with HeLa cells. Our microscopic examination suggested a decrease in the numbers of adherent EPEC upon treatment with a concentration (non-toxic for planktonic EPEC) (30 μM) of vancomycin (**Fig. 6B**). Vancomycin treatment also resulted in lower pedestal formation (**Fig. 6B**). Importantly, no effect of vancomycin (30 μM) could be detected on planktonic EPEC bacteria (**Fig. 6C**). To test the influence of vancomycin on adherent EPEC, we infected HeLa cells with wild type EPEC, and treated them with Vancomycin (30 μM). After infection adherent EPEC were detached from the host using Triton X-100, and plated for colony forming units. The data show that attached EPEC become much more sensitive to vancomycin in comparison to nonattached EPEC. (**Fig. 6D**).

**Fig 6.**
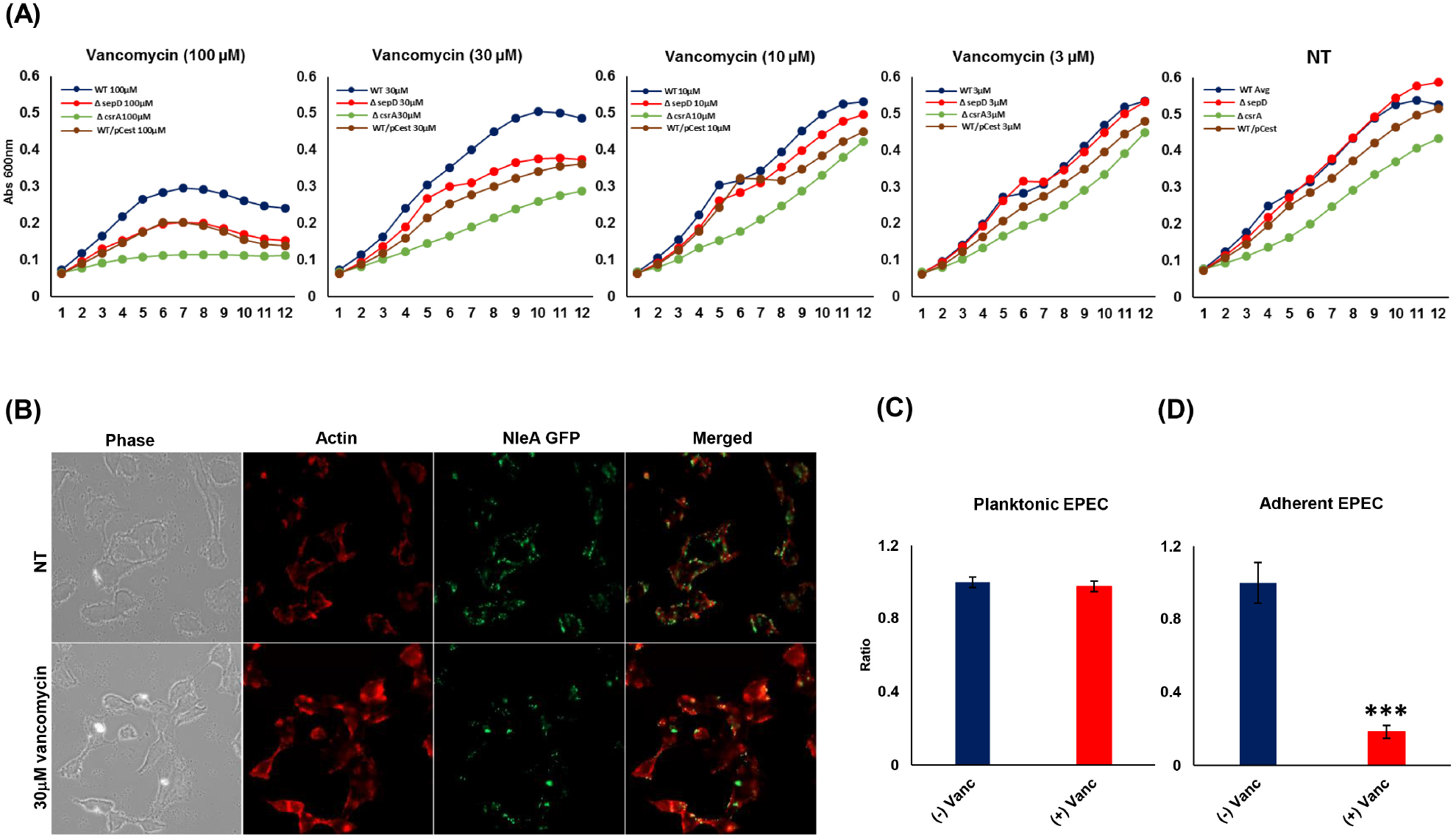
T3SS activation upon attachment to host cells increases the permeability of EPEC. (A) The permeability of EPEC strains was tested in T3SS activated EPEC (Δ*sepD* and WT/pCesT) vs wild type (WT) by a vancomycin resistance assay, where the growth curve was monitored at OD_600_ throughout 12 hours. The vancomycin sensitivity of T3SS activated strains vs wild type (WT) at various concentrations (3, 10, 30, 100 μM) and its respective non-treated (NT) control are shown (A). To confirm the CsrA-mediated response to T3SS activation we also tested the vancomycin permeability of *csrA* mutants. The data is presented as mean ± SEM (N=3). *NleA*-GFP EPEC strain, which exhibits fluorescence upon attachment to the host cells, was examined microscopically. *NleA*-GFP EPEC were grown in DMEM high glucose medium in the presence or absence (non-treated; “NT”) of a concentration (non-toxic for planktonic EPEC) of vancomycin (30 μM) for 4.5 hours in a 24 well plate. The cells were fixed and stained with Rhodamine Phalloidin for actin staining. The infection was visualized using Axio Observer Z1 microscope and image was processed with Zen pro-2012 (B). Planktonic EPEC treated with vancomycin (30 μM), showed no effect on their growth (C). To study the effect of vancomycin on adherent cells, HeLa cells were infected with wild type EPEC in the presence of vancomycin (30 μM). After 3 hours of infection, attached EPEC were recovered and plated on LB agar. Following overnight incubation at 37°C, colonies were counted, and the ratio of vancomycin treated to non-treated was assessed (D). The data shown is an average of 3 experiments performed (n=6 in each experiment) ± SEM.***, p<0.001.

To further define the influence of T3SS on the interactions between the host and bacteria-membrane, we cultured the bacterial strains with diluted serum, and noted a reduced resistance to serum in Δ*sepD* and WT/pCesT mutants (**Fig. 7**).

**Fig S7.**
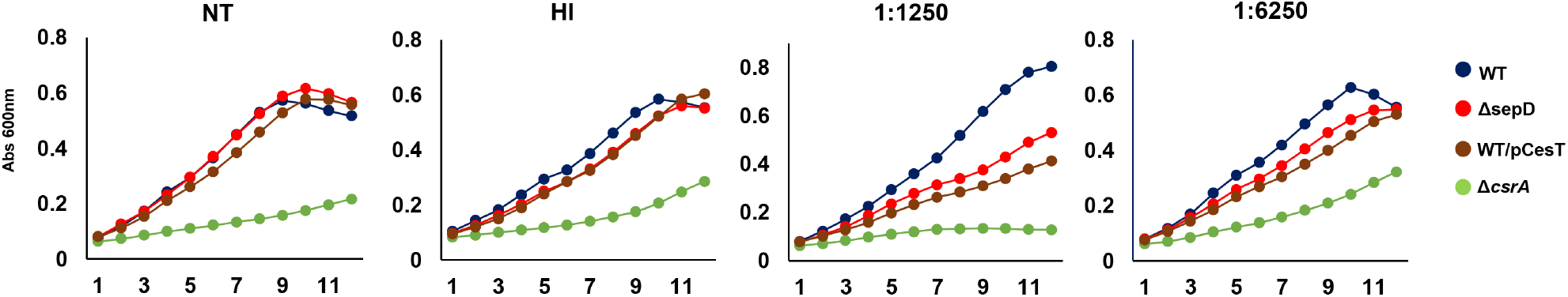
Following T3SS activation EPEC exhibit lower resistance to serum bactericidal components. EPEC strains: wild type, ΔsepD, ΔcsrA and WT/pCesT were grown overnight in media containing human sera of different dilutions, or no serum (non-treated: NT), or heat inactivated sera (HI) at 370 C in a Spark 10M microplate reader. Bacterial growth kinetics was monitored at hourly intervals for 12 hours. The data represented are mean value of N=3 replicates.

## Discussion

In this study, we defined a modulation of membrane lipid composition of pathogenic *E coli* by a secretion system - the T3SS, and demonstrated its implications on membrane function.

An important motivation for carrying out this study was elucidating how attachment to the host and the activation of the T3SS would influence EPEC membrane composition and metabolism. We also expanded the scope of the metabolic regulation by CsrA. Many studies demonstrate the impact of CsrA on *E. coli* central carbon metabolism and physiology, mostly in the laboratory K12 model strains (22, 26-29), but also in other *E. coli* isolates (41), including EPEC (31). Our study elucidates a new aspect of the CsrA function that was previously overlooked. We show here the critical impact of T3SS/CesT regulation of CsrA in controlling *E. coli* lipid metabolism and membrane function. Linking the changes, we found in lipid metabolism to changes in other metabolic networks, and in particular to central carbon metabolism, would be of interest. Importantly, our results point to the dependency of EPEC lipid metabolism on host attachment, and associate it with CsrA repression. Most notably, a shift from PLs to LysoPLs and a shift from menaquinones and ubiquinones towards the production of undecaprenyl lipids. We consider that EPEC Δ*csrA* mutant would only partially phenocopy the physiological state of attached EPEC. This is since this mutant shows reduced growth rate, and in addition may collect suppressor mutations. To better mimic T3SS activation upon host attachment, we capitalized on engineered EPEC strains that transiently and partially inhibit CsrA (as in attached EPEC) and show a growth rate similar to that of the wild type EPEC in our experimental conditions. Using these strains, we carried out a lipidomics study. Altogether, our data is in agreement with available transcript databases of *csrA* mutant EPEC. In a previous analysis by Berndt and colleagues EPEC were cultivated on Modified M9 medium supplement with low glucose (0.02-0.2% glucose (w/v) incubated at 37^?^C, 160 rpm), and their metabolome examined (31). Our lipidomics analyses point to higher levels of LysoPLs in T3SS activated EPEC, whereas the data of Berndt *et al*. disagree with this premise. This discrepancy may reflect differences in experimental conditions used in the two studies. For example, we grew the bacteria statically in DMEM to OD_600_ 0.6, while Brendt *et al*. used modified M9 medium and grew the bacteria to OD_600_ 1.0. Notably, the critical factor in selecting our growth conditions was mimicking infection conditions by keeping them static. The seeming discrepancy could also be related to differences in the methods used for sample preparation. As such, we used a lipid extraction system, whereas Berndt and colleagues used an extraction system optimized for species with medium polarity. Further to the PL remodeling, our unbiased lipidomics analysis suggested alterations in the metabolism of terpenoid-quinones. To better understand the changes in EPEC metabolism following T3SS activation, we mined a database of gene expression alterations in *csrA* EPEC mutant (31). The data mined implied to an upregulation of the undecaprenyl branch of the terpenoid quinone pathway in the *csrA* mutant (**Fig. S6**). Whether and how the observed shifts in lipid metabolism contribute to host colonization are key questions that remain to be addressed in future studies. Moreover, following up on our findings in bacteria *in vivo* will be important to establish their possible clinical implications. One possible mechanism to such potential contribution is the regulation of Zeta potential, reflecting the degree of electrostatic repulsion. Our finding of higher Zeta potentials in the strains that mimic a post-attachment condition may imply that bacterial cells remodel the membrane charge to avert membranous fusion with the host cells.

Bacteria use terpenoids like undecaprenyl phosphate as lipid carriers to assemble numerous glycan polymers that comprise the bacterial envelope. In *E. coli*, undecaprenyl phosphate is required for synthesizing peptidoglycan, O-antigen, and exopolysaccharides (42). Importantly, *idi* removal suppressed a mutation in undecaprenyl pyrophosphate synthase (important for synthesizing undecaprenyl phosphate) (43). Accordingly, our findings that the expression of both *idi* and *ispU* is higher in the *sepD* mutant and in the over-expressing CesT strains, suggest that EPEC attachment to the host cell may remodel its envelope structure. This notion is supported by the considerable modulation of O-antigen content in the envelope of *sepD* and the over-expressing CesT strains. The regulation of envelope structure by T3SS results in an increase in membrane permeability upon the activation of the T3SS, evident from the higher sensitivity of T3SS mutants to large-scaffold antibiotics as vancomycin. The high sensitivity of adherent wild type EPEC to vancomycin underscores the physiological implications of this finding. The dramatic reduction in the numbers of attached EPEC due to concentrations (non-toxic for planktonic EPEC) of vancomycin may also be the result of lower adherence in sub-toxic concentrations of vancomycin. Our lipidomics analyses that point to a considerable increase in the abundance of LysoPLs, under conditions that mimic EPEC attachment are of interest. LysoPLs produced by host intestinal cells induce a cAMP-dependent signaling pathway in infecting *Salmonella*, resulting in the production and secretion of active flagellin (44). Thus, it would be intriguing to test if bacteria might use self-produced LysoPLs as a second messenger. EPEC may also produce LysoPAs to manipulate host signaling, since LysoPA is a potent agonist of a family of five G protein-coupled receptors (GPCRs) associated with G_i_, G_12/13_, G_q,_ and G_s_, which activate a plethora of downstream signaling pathways including PI3K, Ras, Rho, PLC and adenylate cyclase (45-50). Moreover, the interaction of LysoPAs with GPCRs inhibits pro-inflammatory responses induced by lipopolysaccharide (LPS) (46-48, 51). Given the above, it is possible that LysoPL and LysoPA production and secretion may contribute to the capacity of EPEC to evade the host immune response.

In conclusion, our work provides a comprehensive account of the T3SS-dependent lipid metabolism and membrane biogenesis via CsrA repression in EPEC. They provides the foundation for the study of the regulation of lipid metabolism and membrane function by bacterial secretion systems. Our data provide a first insight into the remodeling of bacterial membrane lipid composition upon host attachment, suggesting a metabolic switch from planktonic to the cell-adherent lifestyle.

## Methods

### Bacteria cultures

Bacterial strains and plasmids used are presented in **Table S2** and primers in **Table S3**. All bacterial strains are isogenic. Bacteria were grown in Luria-Bertani (LB) broth supplemented, when needed for subculture, with ampicillin (100 µg/mL), streptomycin (50 µg/mL), chloramphenicol (25 µg/mL), kanamycin (30 µg/mL), or tetracycline (10 µg/mL). For lipid extraction EPEC were grown statically at 37°C overnight in LB medium with respective antibiotics. For all experiments, overnight grown bacterial cultures were then diluted 1:50 using high glucose Dulbecco’s modified Eagle medium (DMEM; Biological Industries) lacking pyruvate and glutamate, and statically grown at 37°C. They were kept in these growth conditions until reaching OD_600_ 0.6.

To express CesT in strains containing the pCesT plasmid, 0.05 mM IPTG was added 3 hours after culturing in DMEM. Bacterial growth was extended to 0.6 Abs at OD_600_. To express CesT in strains containing the pCesT plasmid, 0.05 mM IPTG was added 3 hours after culturing in DMEM Bacterial growth was extended to 0.6 Abs at OD_600_.

### Cloning, GFP fusion, and bacterial strain construction

For GFP fusion, bacteria were electroporated with pKD46 plasmid harboring λ Red genes (γ, β, and Exo) (52). The tet-sacB cassette (53) was introduced downstream to the desired gene of interest and replaced with GFP insertion. The primer sequence for GFP fusions is given as **Table S3**. GFP fusions were verified using PCR with flanking primers. Primers for PCR confirmation of GFP fusions are shown in **Table S3**. For overexpression of CesT levels, the pKD46 plasmids were cured, and pSA10 plasmid containing *CesT* was electroporated. Overexpression of CesT was induced upon IPTG induction.

### RNA extraction and reverse transcription PCR

Overnight grown EPEC were subcultured in DMEM high glucose medium. Following centrifugation, cell pellets were reconstituted in TAE buffer and three freeze thaw cycles were performed. Total RNA was extracted (Zymo kit Direct-zol™ RNA Miniprep) and RNA (1.5μg) was treated with RQ1 Dnase I (Promega; 1 U/µg RNA). cDNA was synthesized with a qPCRBIO high-quality cDNA synthesis kit, and quantified by real-time PCR using a SYBR green mix (Absolute SYBR Green ROX mix; Thermo). 16S rRNA (rrsB) was used as house keeping gene. Primers for qRT-PCR are shown in **Table S3**.

#### Solvents for lipidomics analysis

Acetonitrile, methanol (both ultra-LC-MS grade), chloroform, and water (HPLC-MS grade) were supplied by JT Baker; isopropanol (HPLC-MS grade) from ChemSolute; and formic acid (HPLC-MS grade) by TCI. Ammonium fluoride (>99%) was supplied by Sigma-Aldrich. Internal standard mix – EquiSPLASH LIPIDOMIX (MS-quantitative grade) was obtained from Avanti Polar Lipids.

#### Sample preparation for lipidomics analyses

Cells were centrifuged (5000 g for 10 minutes at 4°C), washed twice with 20 mL of cold phosphate-buffered saline (PBS), and bacterial pellets (45mg, wet weight) were immediately snap-frozen in liquid nitrogen and stored for overnight at -80°C and analyzed the following day. Bacteria (45 mg, wet pellet) were reconstituted in 400 µL of LC-MS grade water and transferred to glass tubes. Following the addition of 800 µL of ice-cold methanol and internal standards (EquiSPLASH LIPIDOMIX), samples went through five cycles of 30-sec ultra-sonication at 4°C for quenching and complete lysis. 400 µL of cold chloroform was added, and another sequence of 30-sec x 5 sonication cycles at 4°C was carried out. The tubes were incubated at room temperature for 30 minutes with occasional mixing (54) and then centrifuged at 770 g for 10 minutes at 4°C for phase separation. The lower chloroform phase was transferred to clean glass tubes, and the protein disk at interface was re-extracted with the same solvent system. Protein disk extracts were then pooled with the respective sample. Samples were concentrated in vacuum concentrator (126 SC210A SpeedVac; Thermo Scientific) and reconstituted in 200 µL 95 % acetonitrile, 0.1 FA. Samples were filtered through Acrodisc^®^ PTFE membrane filters 0.2µm (Pall Corporation, USA) and transferred to Waters^®^ACQUITY UPLC 700µl round 96 well sample plate.

### UPLC-MS analysis

LC-MS analysis was carried out in a Waters Acquity UPLC H-Class (Waters, Milford, MA, USA) and Xevo G2-XS Q-ToF-High resolution, High Mass Accuracy Q-ToF (Waters, Manchester, UK) system. LC-MS runs for lipidomics analyses were performed using a UPLC CSH C18 column (100 mm × 2.1 mm, 1.7 μm; Waters). The column temperature was maintained at 60^°^C. The mobile phase consisted of 0.1% (vol/vol) FA in water (A), 0.1% FA in acetonitrile (B), and isopropanol (C). A flow rate of 0.4 mL/minute was used with a linear gradient (**Table S4**). ESI-MS was calibrated using sodium formate, and leucine enkephalin was used as the lock mass (m/z 556.2771, 200 pg/mL) and continuously infused at 6 µL/min. The capillary spray was maintained at 3.0 kV; the data were acquired in positive and negative mode with collision energies of 15-45 eV and 30-60 eV, respectively. Full-scan and MS^E^ data acquisition were performed, ranging from 30-2000 Dalton. Argon was used as the collision gas for collision-induced dissociation. The lock mass was continuously infused at 6 µL/min. Ammonium fluoride was used for post-column derivatization to improve the yields of the neutral charged lipids in the ES+ mode. MassLynx 4.1 (Waters Corporation, Milford, MA, USA) was used to control the instrument, calculate accurate masses, and mass spectral visualization.

### Lipidomics analyses

Progenesis QI (Nonlinear Dynamics, Newcastle, United Kingdom) was used for spectra deconvolution, alignment, and feature identification. Blank samples (solvents that went through the same sample preparation with no bacteria) were used to exclude artifactual mass features. Mass features eluted at t > 1 minute, with minimum intensity higher than 100, with the lowest mean abundance in the blank, and fold change over 100 from blank, were used for analysis. Following quantile normalization, multivariate tests were carried out us Ezinfo 3.0 (Umetrics AB, Umea Sweden) and Metaboanalyst 4.0 (55). Variable Importance in projection analysis was performed and discriminant metabolic features in Δ*sepD* and Δ*csrA* strains were determined. Full-scan and MS^E^ mass spectra were acquired from all masses of 30–2,000 Daltons. Identification of mass features was carried out using 18 metabolite libraries compatible with Progenesis QI, as well as our internal library, based on mass accuracy < 5 ppm, isotope pattern, fragmentation pattern, and elution time.

### Data mining for gene expression

Changes in transcript levels mediated by *csrA* (31) were previously reported. We mined the publicly available datasets that resulted from these studies for network-based analysis of lipid metabolism regulation by T3SS activation and *csrA* inhibition. The pathways of glycerophospholipid and terpenoid quinone pathways were adapted from KEGG database and are presented and modified to allow presentation in Figure. The list of enzymes involved in *E. coli* lipid synthesis was extracted from the KEGG database.

### Determination of GFP fluorescence intensity

The strains with GFP fusion were grown overnight in LB containing chloramphenicol at 37°C. The overnight cultures were subcultured (1:100) in DMEM high glucose medium. IPTG was added after 3 hours, and growth continued for an additional 3 hours. The cells were washed and suspended in PBS. The fluorescence of GFP was measured at 485 nm excitation and 510 nm emission using the Spark 10M microplate reader (Tecan Trading AG, Switzerland) and normalized to their respective optical densities (OD_600_) (25).

### Bacterial cell size and Zeta potential measurements

EPEC strains were subcultured in DMEM at 37°C to an OD_600_ of 0.6. Bacteria were loaded on polylysine-coated glass slides; a coverslip was mounted on top, and the cells were visualized under a phase-contrast Axio Observer Z1 microscope (Zeiss, Germany). System control and image processing were performed with Zen pro-2012 (Zeiss, Germany). Bacterial cell size was measured using Image J software.

Zeta potential was measured as follows: Bacteria were centrifuged for 3 minutes at 16,000 g at 4°C, washed and reconstituted with 1 mL of LCMS grade water. Zeta potential was determined using Zetasizer Nano ZS (Malvern Instruments, UK) at 25°C. Zeta potential was calculated using the Smoluchowski model. Data acquisition from 15 events was recorded for every sample, and the average Zeta potential was determined from four replicates (56).

### Membrane staining

For staining bacterial membranes, static cultures of EPEC strains were subcultured in DMEM at 37°C to an OD_600_ of 0.6. The cells were washed with PBS, treated with 1 mg/mL FM4-64 (Molecular Probes, Invitrogen), and spotted on poly-L-lysine coated coverslips. Bacteria were visualized in an Eclipse Ti microscope (Nikon, Japan), equipped with a CoolSnap HQII camera (Photometrics, Roper Scientific, USA). System control, image processing, and fluorescence intensity measurement were performed with NIS Elements AR 4.3 (Nikon, Japan).

### LPS O-antigen Quantification

For staining bacterial O-antigen, static cultures of EPEC strains were subcultured in DMEM at 37°C to an OD_600_ of 0.6. Bacteria were washed with PBS and spotted onto poly-L-Lysine coated coverslips. Bacteria on the coverslips were then fixed with 2% paraformaldehyde and 0.01% glutaraldehyde in sodium cacodylate buffer (0.1 M, pH 7.2) for 15 minutes at 25^?^C. Subsequently, coverslips were washed 3 times in PBS, incubated for 30 minutes with 2% BSA and then with rabbit anti-O127 antibody (1:500, 2% BSA), for 1 hour at 25^?^C. Coverslips were washed 3 times with PBS and incubated for 1 hour at 25^?^C with Alexa 488 conjugated goat anti-rabbit antibody (1:1000). Coverslips were then washed 3 times with PBS and fixed with 2.5% glutaraldehyde in sodium cacodylate buffer (0.1 M, pH 7.2) for 15 minutes at 25^?^C. Bacteria were visualized by Eclipse Ti microscope (Nikon, Japan), equipped with CoolSnap HQII camera (Photometrics, Roper Scientific, USA). System control, image processing and fluorescence intensity measurement were performed with NIS Elements AR 4.3 (Nikon, Japan).

### LPS Extraction and O-antigen quantification

For LPS extraction overnight grown cultures in Luria Broth (LB), supplemented with antibiotics (see Bacteria cultures) were subcultured in DMEM at 37°C to an OD_600_ of 0.6. The cells were centrifuged at 10,000x g for 10 minutes and the pellet was collected. The pellets were reconstituted in 200 µL Laemelli buffer with bromophenol blue dye, and kept at 95^?^c for 15 minutes and allowed to cool at room temperature for 15 minutes. 10 μL of Proteinase K solution (10 mg/mL) was added to the samples and incubated at 59^?^C for 3 hours. 200 μL of ice cold water saturated phenol was added to the samples. The samples were vortexed for approximately 5 to 10 seconds, and incubated at 65^?^C for 15 minutes with occasional vortexing. After cooling to room temperature, 1 mL of diethyl ether was added to each sample, followed by vortex (5-10 seconds). The samples were centrifuged (13000 g, 10 minutes). The lower blue layer was carefully removed, and samples were run on 12% SDS-polyacrylamide gels (Mini-PROTEAN® TGX Stain-Free™). The bands were transferred to 0.2 µM Nitrocellulose membrane (Trans Blot Turbo, Biorad laboratories, USA). The blot was blocked with BSA (Bovine serum Albumin) and skim milk (0.6%) in TBS overnight at 4^?^C. The membrane was then incubated with Rabbit-AntiO127 antibody (1:1000) for 1 hour at room temperature, washed 3 times with TBST and incubated with Anti Rabbit IgG (Whole molecule)-AP antibody (1:10000) for one hour. The membrane was then washed with TBST two times. Finally, the membrane was washed with AP buffer (100 mM Tris-HCl [pH 9.0], 150 mM NaCl, and 1 mM MgCl_2_). The membrane was visualized using GeldocTm Ezimagin (Biorad USA) with transwhite background.

### Vancomycin permeability assays

Overnight grown EPEC strains were subcultured in DMEM medium in 96 well transparent flatbottom plate. Vancomycin (Sigma Aldrich, Louis, MO, USA; 0, 3, 10, 30, 100 µM) was incubated with EPEC strains at 37^?^C. The growth rate was monitored at 600 nm for 12 hours using Spark 10M microplate reader (Tecan Trading AG, Switzerland).

HeLa cells were seeded in a 24-well plates (Nunc) at a density of 7 ×10^4^ cells per well and grown overnight in DMEM supplemented with 10% fetal calf serum (Biological Industries) and antibiotics (penicillin-streptomycin solution; Biological Industries). They were infected with statically overnight grown wildtype EPEC *nleA gfp* (1:100). The infection was carried out in the presence of vancomycin (30 µM). After 3 hours of infection, the wells were washed with PBS twice. To terminate the infection, the cells were then fixed (3.7% formaldehyde in PBS), washed, perforated (PBS, 0.25% Triton X-100 for 10 min), washed, stained with phalloidin rhodamine (Sigma), and analyzed by fluorescence microscopy Axio Observer Z1 microscope (Zeiss, Germany). System control and image processing were performed with Zen pro-2012 (Zeiss, Germany).

HeLa cells were seeded in a 24-well plates (Nunc) at a density of 7 ×10^4^ cells per well and grown overnight in DMEM supplemented with 10% fetal calf serum (Biological Industries) and antibiotics (penicillin-streptomycin solution; Biological Industries). HeLa cells were infected with statically overnight grown wild type EPEC (1:100). The infection was carried out in the presence of vancomycin (30 µM). After 3 hours of infection, the wells were washed with PBS twice. 0.5% of Triton X-100 was added and incubated for 10min at room temperature. Bacteria regained from cell surface were plated on the LB agar and incubated at 37°C overnight. The colonies were counted and the ratio of adherent cells to planktonic cell state was determined.

### Serum bactericidal assay

Overnight grown EPEC strains were subcultured in DMEM medium in 96 well transparent flatbottom plate. Human serum samples were heat inactivated at 56^?^C. Serum samples were diluted from 1:1250-1:6250 and incubated with EPEC strains at 37^?^C. The growth rate was monitored at 600 nm for 12 hours using Spark 10 M microplate reader (Tecan Trading AG, Switzerland).

## Competing interests

We have no competing interests to report.

## Legends for Supplemental Figure and Tables

**Fig S1** Schematic representation of the lipid classes modified by T3SS activation in EPEC. (A) Glycerophopholipid structure representing glycerol unit attached to the two fatty acyl chains (R1 and R2) or single acyl chain (LysoPL). A phosphate head group determines the lipid class, and is represented by X. (X-i) Phosphatidylethanolamine (PE); (X-ii) Phosphatidylglycerol (PG); (X-iii) Phosphatidyl serine. (B) Cardiolipin (CL) is a phospholipid with two phosphatidic acids linked to the two carbons of glycerol unit, hence four acyl chains R1,R2 and R3,R4 attached to the respective glycerol units. (C) Monomeric isoprene unit with five carbons. These isoprenes form different number of repeats represented as n, and may bind to: (Y-i) Di phosphate; (Y-ii) Ubiquinone; (Y-iii) Menaquinone.

**Fig S2** Growth curve of EPEC wild type, ΔsepD, ΔcsrA, ΔsepDΔcesT, and WT/pCesT strains. EPEC strains were grown overnight in LB broth and then cultured in DMEM high glucose medium (lacking glutamine and pyruvate) at 37°C under static conditions, until reaching optical density of 600 nm (OD)_600_ 0.6, at which point bacterial growth was determined.

**Fig S3** Membrane stain analysis showing the lower uptake of membrane stain FM 4-64 in strains with high levels of CesT. Phase-contrast and fluorescent images of EPEC wild type (WT) and mutants following FM 4-64 membrane staining. Scale bar indicates 5 μm (A), The fluorescence emission from ∼ 10,000 bacteria for each strain, in 3 technical replicates were recorded and the data was presented as mean ± SE. ***, p<0.001 (B).

**Fig S4** Data mining of csrA mutant transcriptome corroborated a csrA-mediated regulation of the expression of phospholipid pathway genes by the T3SS. A metabolic map of phospholipids with data from our lipidomics analysis was integrated with transcriptomic data taken from Berndt et al., 2019 (31) (A). Glycerophospholipid pathway was adapted from Kyoto Encyclopedia of Genes and Genomes (KEGG) for Escherichia coli O127:H6 E2348/69 (EPEC). Lipid classes are represented by circles, with their names in white quadrants. Enzymes are represented by arrows, with their names in colored quadrants. Colors of circles (lipid classes) and quadrants (gene expression) are given as ln2 (fold change), relative to the wild type mean value. Abbreviations used for lipid species names: 13P, Glycerone phosphate; G3P, Glycerol-3-phosphate; AGP, Acylglycerol-3-phosphate; DAGP, 1,2-Diacylglycerol 3-phosphate; CDPDG, 1,2-Diacylglycerol-cytidine 5-diphosphate; GPE, Glycerophosphoethanolamine; LPE, Lysophosophoethanolamine; Lyso PS, Lysophosphoserine; PGP, Phosphatidylglycerophosphate; PE, Phosphatidylethanolamine; PS, Phosphatidylserine; PG, Phosphatidylglycerol. Enzymes are represented by arrows, with their numbers in quadrants. Enzymes common nomenclature is used in the scheme. For an unambiguous identification, enzyme nomenclature (EC number system) is hereby given, along with further commonly used names: [1.1.1.94]- gpsA, glycerol-3-phosphate dehydrogenase; [1.1.5.3]- glpD, sn-glycerol-3-phosphate dehydrogenase; [2.3.1.15]- plsB, glycerol-3-phosphate O-acyltransferase; [2.3.1.274]- plsX, phosphate acyltransferase; [2.3.1.275]- plsY (ygiH), glycerol-3-phosphate acyltransferase; [2.3.1.51]- plsC (parF), 1-acyl-sn-glycerol-3-phosphate acyltransferase; [2.7.1.107]- dgkA, diacylglycerol kinase; [3.6.1.26]- cdh, CDP-diacylglycerol phosphotidylhydrolase, [2.7.7.41]- cdsA, (cds) CDP-diglyceride synthase; [2.7.8.5]- pgsA, phosphatidylglycerophosphate synthetase; [3.1.3.27]- pgpA (yajN), phosphatidylglycerophosphatase A; [2.7.8.8]- pssA (pss), phosphatidylserine synthase; [3.1.1.32]- pldA, outer membrane phospholipase A; [<u>4.1.1.65</u>]- psd, phosphatidylserine decarboxylase; [3.1.1.4]- pldA, outer membrane phospholipase A; [2.3.1.40]- aas, 2-Acyl-sn- glycero-3-phosphoethanolamine O-acyltransferase; [3.1.1.5]- pldB, lysophospholipase L; [3.1.4.46]- glpQ (ugpQ), glycerophosphodiester phosphodiesterase. The T3SS-related shift in the phospholipid composition is mediated by csrA (B-D). The total abundance of identified phospholipids- phosphatidylglycerols (PGs; B) to phosphatidylethanolamines (PEs; C), lysophospholipids (LysoPLs; D) is presented. The expression of key enzymes responsible for the conversion of PLs to LysoPLs was evaluated by RT-qPCR: pldA (E) and ygiH (F). The data is presented as mean ± SE (N=4). **, p<0.01; ***, p<0.001.

**Fig S5** No change noted in the expression of UbiC, MenF and WrbA following CesT accumulation. Quinone terpenoid biosynthesis genes were GFP labeled in EPEC null for CesT or overexpressing it. GFP fluorescence intensity was then measured. Data are presented as mean ± SE (N=3).

**Fig S6** Data mining in the ΔcsrA transcriptome (31) suggests upregulation of undecaprenyl biosynthesis. Network was adapted from Kyoto Encyclopedia of Genes and Genomes (KEGG) for Escherichia coli O127:H6 E2348/69 (EPEC), and modified to include undecaprenyl, ubiquinone and menaquinone branches of terpenoid pathway. Lipids are represented by circles, with their names in colored quadrants. Enzyme are represented by arrows, with their names in colored quadrents. The color of quadrants represent the log2 fold change between the csrA mutant and the wild type (WT) strain. Lipids abbreviations: H6P, 1-Hydroxy-2-methyl-2-butenyl-4-diphosphate; DMA, Dimethylallyl diphosphate; IPE, Isopentenyl diphosphate; GPP, Geranyl diphosphate; FPP, Farnesyl diphosphate; UPP, Undecaprenyl diphosphate; OTP, Octaprenyldiphosphate; PHB, Hydroxybenzoic acid; ISJ, Chorismate; O4HPZ, 4-Hydroxy-3-polyprenyl benzoate; 2OPPP, 2-Octaprenyl phenol; 2O6H, 2-Octaprenyl-6-hydroxyphenol; 2OPMP, 2-Octaprenyl-6-methoxy phenol; 2OPMB, 2-Octaprenyl-6-methoxy-1,4-benzoquinone; 2OPMMB, 2-Octaprenyl-3-methyl-6-methoxy-1,4-benzoquinone; 2OMHMB, 2-Octaprenyl-3-methyl-5-hydroxy-6-methoxy-1,4-benzoquinone; Q, Ubiquinone; ISC, Isochorismate; SEPHCHC, 2-Succinyl-5-enolpyruvyl-6-hydroxy-3-cyclohexene-1-carboxylate; SHCHC, 6-Hydroxy-2-succinylcyclohexa-2,4-diene-1-carboxylate; OSB, 2-Succinylbenzoate; OSB-COA, 2-Succinylbenzoyl-CoA; DHNA-CoA, 1,4-Dihydroxy-2-naphthoyl-CoA; DHNA, 1,4-Dihydroxy-2-naphthoate; DMKH2, Demethylmenaquinol; MKH2, Menaquinol; MK, Menaquinone. Enzymes common nomenclature is used in the scheme. For an unambiguous identification, enzyme nomenclature (EC number system) is hereby given, along with further commonly used names: [<u>1.17.7.4</u>]- ispH (yaaE, lytB), 4-hydroxy-3-methylbut-2-enyl diphosphate reductase; [2.5.1.1, 2.5.1.10]- ispA, farnesyl diphosphate synthase; [2.5.1.90]- ispB (cel, yhbD), all-trans-octaprenyl- diphosphate synthase; [2.5.1.31]- ispU (uppS, rth, yaeS), ditrans,polycis-undecaprenyl-diphosphate synthase; [4.1.3.40]- ubiC, chorismate lyase; [2.5.1.39] ubiA, 4-hydroxybenzoate octaprenyltransferase; [2.5.1.129]- ubiX (dedF), 3-octaprenyl-4-hydroxybenzoate carboxy-lyase; [<u>1.14.13.240</u>]- ubiI (visC),), 2-octaprenyl phenol 6- hydroxylase; [2.1.1.222, 2.1.1.64]- ubiG (pufX, yfaB), bifunctional 3-demethylubiquinone-9 3-O-methyltransferase and 2-octaprenyl-6-hydroxyphenolmethylase; [1.14.13.-]- ubiH (acd, visB) 2-octaprenyl-6-methoxyphenol 4- hydroxylase; [2.1.1.201]- ubiE (yigO), bifunctional 2-octaprenyl-6-methoxy-1,4-benzoquinone methylase and demethylmenaquinone methyltransferase; [1.14.99.60]- ubiF (yleB), 2-octaprenyl-3-methyl-6-methoxy-1,4- benzoquinol oxygenase, 2-octaprenyl-3-methyl-6-methoxy-1,4-benzoquinol hydroxylase; [2.1.1.222, 2.1.1.64]- ubiG (pufX, yfaB) bifunctional 3-demethylubiquinone-9 3-O-methyltransferase and 2-octaprenyl-6-hydroxyphenol methylase; [5.4.4.2]- menF (yfbA), isochorismate synthase 2; [2.2.1.9]- menD, 2-succinyl-5-enolpyruvyl-6-hydroxy-3- cyclohexene-1-carboxylatesynthase; [<u>4.2.99.20</u>]- menH (yfbB), 2-succinyl-6-hydroxy-2,4-cyclohexadiene-1- carboxylate synthase; [4.2.1.113]- menC, o-succinylbenzoate synthase; [6.2.1.26]- menE, o-succinylbenzoateCoA ligase; [4.1.3.36]- menB, 1,4-dihydroxy-2-naphthoyl-CoA synthase; [3.1.2.28]- ydiI (menI), 1,4-dihydroxy-2- naphthoyl-CoA hydrolase; [2.5.1.74]- menA (yiiW), 1,4-dihydroxy-2-naphthoate octaprenyltransferase; [2.1.1.163, 2.1.1.201]- ubiE (menG, yigO), demethylmenaquinone methyltransferase /2-methoxy-6-octaprenyl-1,4- benzoquinol methylase; [1.6.5.2]- wrbA(ytfG), quinoneoxidoreductase.

**Table S1** Identified lipids (taken from the Lipidomics analysis of Δ*sepD* and WT/pCesT EPEC with respect to wild-type) PG, Phosphatidyl glycerol; PE, phosphatidyl ethanolamide; PS, phosphatidyl serine; PA, phosphatidic acid; CL, cardiolipin.The lipid species were identified by MS^E^. The Data shown in the table represent the fold change calculated from the abundance of each lipid in the Δ*sepD* and WT/pCesT from the wild type.

*, an intermediate ubiquinones.

**Table S2** Strains and plasmids used for cloning

**Table S3** Primer sequence for GFP fusions in EPEC

**Table S4** The linear gradient for the lipidomics analyses

## Notes

### Competing Interest Statement

The authors have declared no competing interest.

## References

1. Duong F, Eichler J, Price A, Leonard MR, Wickner W. 1997. Biogenesis of the gram-negative bacterial envelope. Cell 91:567–73.

2. Lundstedt E, Kahne D, Ruiz N. 2021. Assembly and Maintenance of Lipids at the Bacterial Outer Membrane. Chem Rev 121:5098–5123.

3. Deng W, Marshall NC, Rowland JL, McCoy JM, Worrall LJ, Santos AS, Strynadka NCJ, Finlay BB. 2017. Assembly, structure, function and regulation of type III secretion systems. Nat Rev Microbiol 15:323–337.

4. Nataro JP, Kaper JB. 1998. Diarrheagenic Escherichia coli. Clin Microbiol Rev 11:142–201.

5. Elliott SJ, Wainwright LA, McDaniel TK, Jarvis KG, Deng YK, Lai LC, McNamara BP, Donnenberg MS, Kaper JB. 1998. The complete sequence of the locus of enterocyte effacement (LEE) from enteropathogenic Escherichia coli E2348/69. Mol Microbiol 28:1–4.

6. Garmendia J, Frankel G, Crepin VF. 2005. Enteropathogenic and enterohemorrhagic Escherichia coli infections: translocation, translocation, translocation. Infect Immun 73:2573–85.

7. Gaytan MO, Martinez-Santos VI, Soto E, Gonzalez-Pedrajo B. 2016. Type Three Secretion System in Attaching and Effacing Pathogens. Front Cell Infect Microbiol 6:129.

8. Mellies JL, Elliott SJ, Sperandio V, Donnenberg MS, Kaper JB. 1999. The Per regulon of enteropathogenic Escherichia coli : identification of a regulatory cascade and a novel transcriptional activator, the locus of enterocyte effacement (LEE)-encoded regulator (Ler). Mol Microbiol 33:296–306.

9. Sanchez-SanMartin C, Bustamante VH, Calva E, Puente JL. 2001. Transcriptional regulation of the orf19 gene and the tir-cesT-eae operon of enteropathogenic Escherichia coli. J Bacteriol 183:2823–33.

10. Kenny B, DeVinney R, Stein M, Reinscheid DJ, Frey EA, Finlay BB. 1997. Enteropathogenic E. coli (EPEC) transfers its receptor for intimate adherence into mammalian cells. Cell 91:511–20.

11. Mills E, Baruch K, Aviv G, Nitzan M, Rosenshine I. 2013. Dynamics of the type III secretion system activity of enteropathogenic Escherichia coli. mBio 4.

12. Little DJ, Coombes BK. 2018. Molecular basis for CesT recognition of type III secretion effectors in enteropathogenic Escherichia coli. PLoS Pathog 14:e1007224.

13. Runte CS, Jain U, Getz LJ, Secord S, Kuwae A, Abe A, LeBlanc JJ, Stadnyk AW, Kaper JB, Hansen AM, Thomas NA. 2018. Tandem tyrosine phosphosites in the Enteropathogenic Escherichia coli chaperone CesT are required for differential type III effector translocation and virulence. Mol Microbiol 108:536–550.

14. Lai Y, Rosenshine I, Leong JM, Frankel G. 2013. Intimate host attachment: enteropathogenic and enterohaemorrhagic Escherichia coli. Cell Microbiol 15:1796–808.

15. Login FH, Jensen HH, Pedersen GA, Amieva MR, Nejsum LN. 2018. The soluble extracellular domain of E-cadherin interferes with EPEC adherence via interaction with the Tir:intimin complex. FASEB J doi:10.1096/fj.201800651:fj201800651.

16. Katsowich N, Elbaz N, Pal RR, Mills E, Kobi S, Kahan T, Rosenshine I. 2017. Host cell attachment elicits posttranscriptional regulation in infecting enteropathogenic bacteria. Science 355:735–739.

17. Mills E, Baruch K, Charpentier X, Kobi S, Rosenshine I. 2008. Real-time analysis of effector translocation by the type III secretion system of enteropathogenic Escherichia coli. Cell Host Microbe 3:104–13.

18. Goddard PJ, Sanchez-Garrido J, Slater SL, Kalyan M, Ruano-Gallego D, Marches O, Fernandez LA, Frankel G, Shenoy AR. 2019. Enteropathogenic Escherichia coli Stimulates Effector-Driven Rapid Caspase-4 Activation in Human Macrophages. Cell Rep 27:1008–1017 e6.

19. Diaz-Guerrero M, Gaytan MO, Soto E, Espinosa N, Garcia-Gomez E, Marcos-Vilchis A, Andrade A, Gonzalez-Pedrajo B. 2021. CesL Regulates Type III Secretion Substrate Specificity of the Enteropathogenic E. coli Injectisome. Microorganisms 9.

20. Portaliou AG, Tsolis KC, Loos MS, Balabanidou V, Rayo J, Tsirigotaki A, Crepin VF, Frankel G, Kalodimos CG, Karamanou S, Economou A. 2017. Hierarchical protein targeting and secretion is controlled by an affinity switch in the type III secretion system of enteropathogenic Escherichia coli. EMBO J 36:3517–3531.

21. Ye F, Yang FL, Yu RJ, Lin X, Qi JX, Chen ZJ, Cao Y, Wei YQ, Gao GF, Lu GW. 2018. Molecular basis of binding between the global post-transcriptional regulator CsrA and the T3SS chaperone CesT. Nature Communications 9.

22. Potts AH, Vakulskas CA, Pannuri A, Yakhnin H, Babitzke P, Romeo T. 2017. Global role of the bacterial post-transcriptional regulator CsrA revealed by integrated transcriptomics. Nat Commun 8:1596.

23. Romeo T, Gong M, Liu MY, Brun-Zinkernagel AM. 1993. Identification and molecular characterization of csrA, a pleiotropic gene from Escherichia coli that affects glycogen biosynthesis, gluconeogenesis, cell size, and surface properties. J Bacteriol 175:4744–55.

24. Ye F, Yang F, Yu R, Lin X, Qi J, Chen Z, Cao Y, Wei Y, Gao GF, Lu G. 2018. Molecular basis of binding between the global post-transcriptional regulator CsrA and the T3SS chaperone CesT. Nat Commun 9:1196.

25. Elbaz N, Socol Y, Katsowich N, Rosenshine I. 2019. Control of Type III Secretion System Effector/Chaperone Ratio Fosters Pathogen Adaptation to Host-Adherent Lifestyle. mBio 10.

26. Sowa SW, Gelderman G, Leistra AN, Buvanendiran A, Lipp S, Pitaktong A, Vakulskas CA, Romeo T, Baldea M, Contreras LM. 2017. Integrative FourD omics approach profiles the target network of the carbon storage regulatory system. Nucleic Acids Res 45:1673–1686.

27. McKee AE, Rutherford BJ, Chivian DC, Baidoo EK, Juminaga D, Kuo D, Benke PI, Dietrich JA, Ma SM, Arkin AP, Petzold CJ, Adams PD, Keasling JD, Chhabra SR. 2012. Manipulation of the carbon storage regulator system for metabolite remodeling and biofuel production in Escherichia coli. Microb Cell Fact 11:79.

28. Sabnis NA, Yang H, Romeo T. 1995. Pleiotropic regulation of central carbohydrate metabolism in Escherichia coli via the gene csrA. J Biol Chem 270:29096–104.

29. Suzuki K, Wang X, Weilbacher T, Pernestig AK, Melefors O, Georgellis D, Babitzke P, Romeo T. 2002. Regulatory circuitry of the CsrA/CsrB and BarA/UvrY systems of Escherichia coli. J Bacteriol 184:5130–40.

30. Deng W, Yu HB, Li Y, Finlay BB. 2015. SepD/SepL-dependent secretion signals of the type III secretion system translocator proteins in enteropathogenic Escherichia coli. J Bacteriol 197:1263–75.

31. Berndt V, Beckstette M, Volk M, Dersch P, Bronstrup M. 2019. Metabolome and transcriptome-wide effects of the carbon storage regulator A in enteropathogenic Escherichia coli. Sci Rep 9:138.

32. Morin M, Ropers D, Letisse F, Laguerre S, Portais JC, Cocaign-Bousquet M, Enjalbert B. 2016. The post-transcriptional regulatory system CSR controls the balance of metabolic pools in upper glycolysis of Escherichia coli. Mol Microbiol 100:686–700.

33. Yoshimura M, Oshima T, Ogasawara N. 2007. Involvement of the YneS/YgiH and PlsX proteins in phospholipid biosynthesis in both Bacillus subtilis and Escherichia coli. BMC Microbiol 7:69.

34. Wang C, Liwei M, Park JB, Jeong SH, Wei G, Wang Y, Kim SW. 2018. Microbial Platform for Terpenoid Production: Escherichia coli and Yeast. Front Microbiol 9:2460.

35. Wang X, Quinn PJ. 2010. Lipopolysaccharide: Biosynthetic pathway and structure modification. Prog Lipid Res 49:97–107.

36. Touz ET, Mengin-Lecreulx D. 2008. Undecaprenyl Phosphate Synthesis. EcoSal Plus 3.

37. Meganathan R, Kwon O. 2009. Biosynthesis of Menaquinone (Vitamin K2) and Ubiquinone (Coenzyme Q). EcoSal Plus 3.

38. Taniguchi Y, Choi PJ, Li GW, Chen H, Babu M, Hearn J, Emili A, Xie XS. 2010. Quantifying E. coli proteome and transcriptome with single-molecule sensitivity in single cells. Science 329:533–8.

39. Peleg A, Shifrin Y, Ilan O, Nadler-Yona C, Nov S, Koby S, Baruch K, Altuvia S, Elgrably-Weiss M, Abe CM, Knutton S, Saper MA, Rosenshine I. 2005. Identification of an Escherichia coli operon required for formation of the O-antigen capsule. J Bacteriol 187:5259–66.

40. Muheim C, Gotzke H, Eriksson AU, Lindberg S, Lauritsen I, Norholm MHH, Daley DO. 2017. Increasing the permeability of Escherichia coli using MAC13243. Sci Rep 7:17629.

41. Revelles O, Millard P, Nougayrede JP, Dobrindt U, Oswald E, Letisse F, Portais JC. 2013. The carbon storage regulator (Csr) system exerts a nutrient-specific control over central metabolism in Escherichia coli strain Nissle 1917. PLoS One 8:e66386.

42. Jorgenson MA, Young KD. 2016. Interrupting Biosynthesis of O Antigen or the Lipopolysaccharide Core Produces Morphological Defects in Escherichia coli by Sequestering Undecaprenyl Phosphate. J Bacteriol 198:3070–3079.

43. MacCain WJ, Kannan S, Jameel DZ, Troutman JM, Young KD. 2018. A Defective Undecaprenyl Pyrophosphate Synthase Induces Growth and Morphological Defects That Are Suppressed by Mutations in the Isoprenoid Pathway of Escherichia coli. J Bacteriol 200.

44. Subramanian N, Qadri A. 2006. Lysophospholipid sensing triggers secretion of flagellin from pathogenic salmonella. Nat Immunol 7:583–9.

45. Chien HY, Lu CS, Chuang KH, Kao PH, Wu YL. 2015. Attenuation of LPS-induced cyclooxygenase-2 and inducible NO synthase expression by lysophosphatidic acid in macrophages. Innate Immun 21:635–46.

46. Zhao J, He D, Su Y, Berdyshev E, Chun J, Natarajan V, Zhao Y. 2011. Lysophosphatidic acid receptor 1 modulates lipopolysaccharide-induced inflammation in alveolar epithelial cells and murine lungs. Am J Physiol Lung Cell Mol Physiol 301: L547–56.

47. Fan H, Zingarelli B, Harris V, Tempel GE, Halushka PV, Cook JA. 2008. Lysophosphatidic acid inhibits bacterial endotoxin-induced pro-inflammatory response: potential anti-inflammatory signaling pathways. Mol Med 14:422–8.

48. Ciesielska A, Hromada-Judycka A, Ziemlinska E, Kwiatkowska K. 2019. Lysophosphatidic acid up-regulates IL-10 production to inhibit TNF-alpha synthesis in Mvarphis stimulated with LPS. J Leukoc Biol 106:1285–1301.

49. Lin S, Han Y, Jenkin K, Lee SJ, Sasaki M, Klapproth JM, He P, Yun CC. 2018. Lysophosphatidic Acid Receptor 1 Is Important for Intestinal Epithelial Barrier Function and Susceptibility to Colitis. Am J Pathol 188:353–366.

50. Wiedmaier N, Muller S, Koberle M, Manncke B, Krejci J, Autenrieth IB, Bohn E. 2008. Bacteria induce CTGF and CYR61 expression in epithelial cells in a lysophosphatidic acid receptor-dependent manner. Int J Med Microbiol 298:231–43.

51. Tsai HC, Han MH. 2016. Sphingosine-1-Phosphate (S1P) and S1P Signaling Pathway: Therapeutic Targets in Autoimmunity and Inflammation. Drugs 76:1067–79.

52. Datsenko KA, Wanner BL. 2000. One-step inactivation of chromosomal genes in Escherichia coli K-12 using PCR products. Proc Natl Acad Sci U S A 97:6640–5.

53. Li XT, Thomason LC, Sawitzke JA, Costantino N, Court DL. 2013. Positive and negative selection using the tetA-sacB cassette: recombineering and P1 transduction in Escherichia coli. Nucleic Acids Res 41:e204.

54. Rowlett VW, Mallampalli V, Karlstaedt A, Dowhan W, Taegtmeyer H, Margolin W, Vitrac H. 2017. Impact of Membrane Phospholipid Alterations in Escherichia coli on Cellular Function and Bacterial Stress Adaptation. J Bacteriol 199.

55. Chong J, Soufan O, Li C, Caraus I, Li S, Bourque G, Wishart DS, Xia J. 2018. MetaboAnalyst 4.0: towards more transparent and integrative metabolomics analysis. Nucleic Acids Res 46:W486–W494.

56. Wyness AJ, Paterson DM, Defew EC, Stutter MI, Avery LM. 2018. The role of zeta potential in the adhesion of E. coli to suspended intertidal sediments. Water Res 142:159–166.

