## Supplemental Material for "Activation of the type 3 secretion system of enteropathogenic *E. coli* leads to remodeling of its membrane composition and function"

Supplementary Material

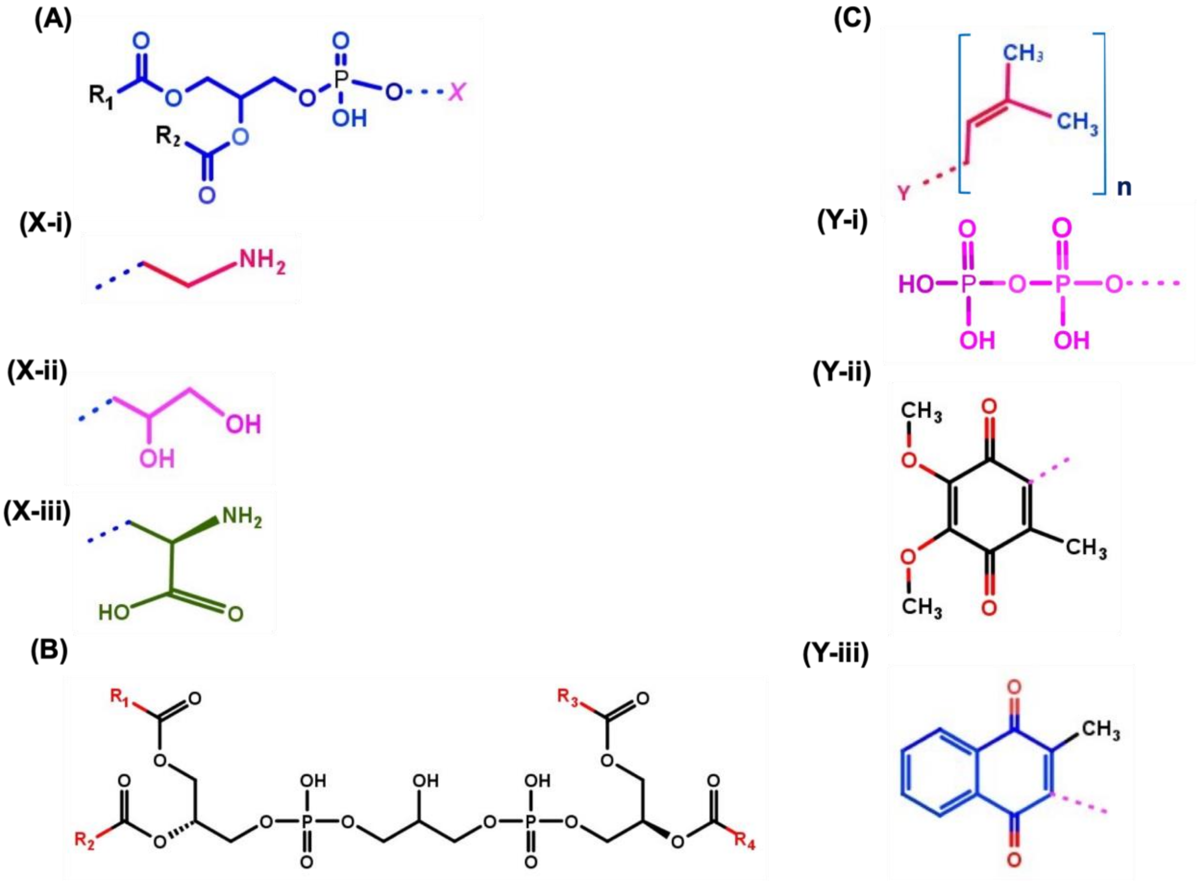


**Fig S1** Schematic representation of the lipid classes modified by T3SS activation in EPEC. (A) Glycerophopholipid structure representing glycerol unit attached to the two fatty acyl chains ($R_1$ and $R_2$) or single acyl chain (LysoPL). A phosphate head group determines the lipid class, and is represented by X. (X-i) Phosphatidylethanolamine (PE); (X-ii) Phosphatidylglycerol (PG); (X-iii) Phosphatidyl serine. (B) Cardiolipin (CL) is a phospholipid with two phosphatidic acids linked to the two carbons of glycerol unit, hence four acyl chains $R_1$,$R_2$ and $R_3$,$R_4$ attached to the respective glycerol units. (C) Monomeric isoprene unit with five carbons. These isoprenes form different number of repeats represented as n, and may bind to: (Y-i) Di phosphate; (Y-ii) Ubiquinone; (Y-iii) Menaquinone.

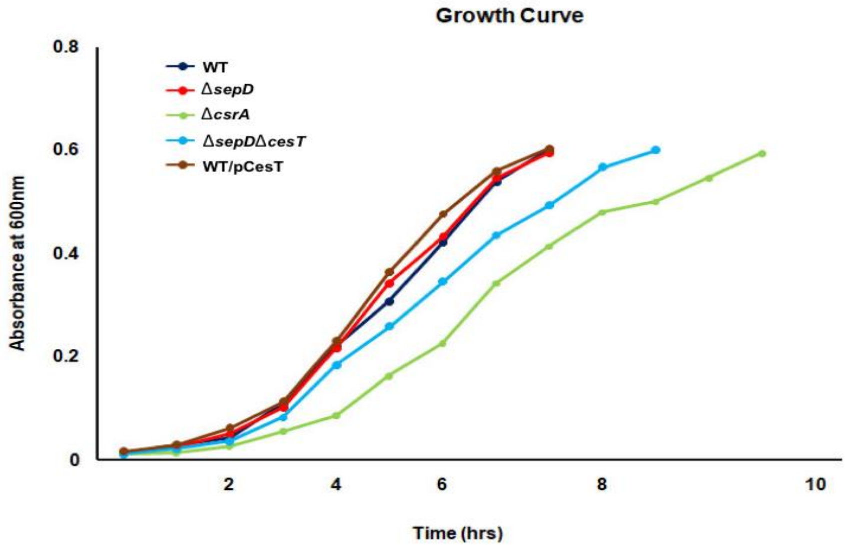


**Fig S2** Growth curve of EPEC wild type, Δ*sepD*, Δ*csrA*, Δ*sepD*Δ*cesT*, and WT/pCesT strains. EPEC strains were grown overnight in LB broth and then cultured in DMEM high glucose medium (lacking glutamine and pyruvate) at 37˚C under static conditions, until reaching optical density of 600 nm $(OD)_{600}$ 0.6, at which point bacterial growth was determined.

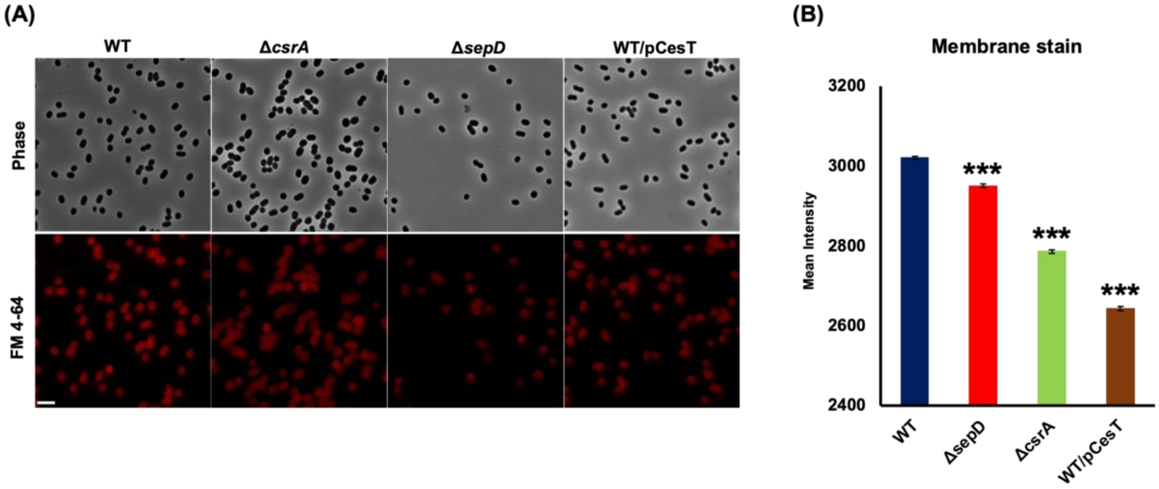


**Fig S3** Membrane stain analysis showing the lower uptake of membrane stain FM 4-64 in strains with high levels of CesT.

Phase-contrast and fluorescent images of EPEC wild type (WT) and mutants following FM 4-64 membrane staining. Scale bar indicates 5 µm (A), The fluorescence emission from ~ 10,000 bacteria for each strain, in 3 technical replicates were recorded and the data was presented as mean ± SE. ***, $p<0.001$ (B).

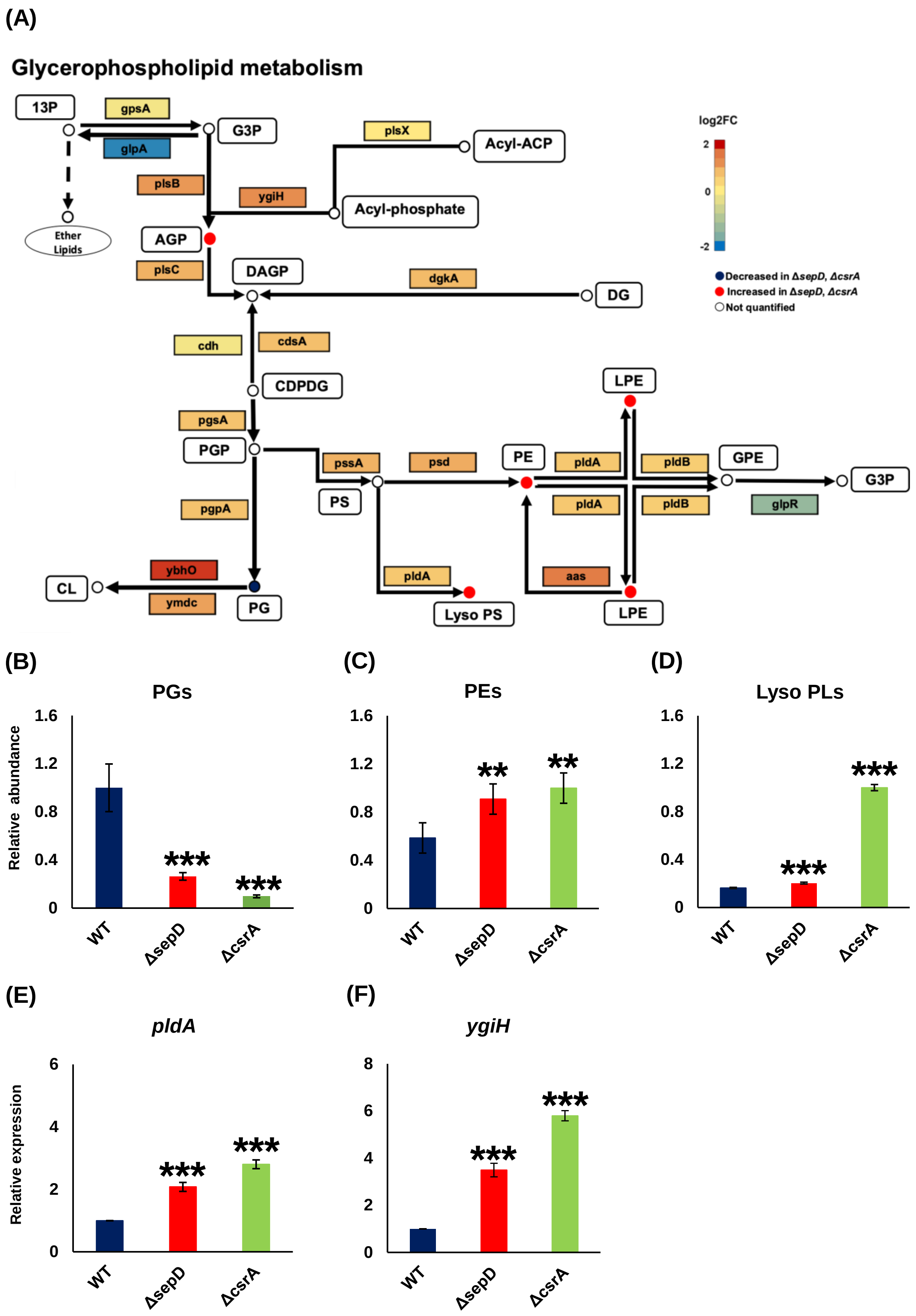

(A)
Glycerophospholipid metabolism
13P
gpsA
glpA
G3P
plsX
Acyl-ACP
plsB
ygiH
Acyl-phosphate
Ether Lipids
AGP
plsC
DAGP
dgkA
DG
cdh
cdsA
CDPDG
pgsA
PGP
pssA
psd
PE
pldA
pldB
GPE
G3P
glpR
LPE
PS
pgpA
ybhO
ymdc
CL
PG
pldA
Lyso PS
aas
LPE
log2FC
2
0
-2
Decreased in ΔsepD, ΔcsrA
Increased in ΔsepD, ΔcsrA
Not quantified
(B)
PGs
Relative abundance
1.6
1.2
0.8
0.4
0
WT
ΔsepD
ΔcsrA
***
***
(C)
PEs
**
**
(D)
Lyso PLs
***
***
(E)
pldA
Relative expression
6
4
2
0
***
***
(F)
ygiH
8
6
4
2
0
***
***

**Fig S4** Data mining of *csrA* mutant transcriptome corroborated a *csrA*-mediated regulation of the expression of phospholipid pathway genes by the T3SS. A metabolic map of phospholipids with data from our lipidomics analysis was integrated with transcriptomic data taken from Berndt *et al*., 2019 (A). Glycerophospholipid pathway was adapted from Kyoto Encyclopedia of Genes and Genomes (KEGG) for *Escherichia coli* O127:H6 E2348/69 (EPEC). Lipid classes are represented by circles, with their names in white quadrants. Enzymes are represented by arrows, with their names in colored quadrants. Colors of circles (lipid classes) and quadrants (gene expression) are given as ln2 (fold change), relative to the wild type mean value. Abbreviations used for lipid species names: 13P, Glycerone phosphate; G3P, Glycerol-3-phosphate; AGP, Acylglycerol-3-phosphate; DAGP, 1,2-Diacylglycerol 3-phosphate; CDPDG, 1,2-Diacylglycerol-cytidine 5-diphosphate; GPE, Glycerophosphoethanolamine; LPE, Lysophosophoethanolamine; Lyso PS, Lysophosphoserine; PGP, Phosphatidylglycerophosphate; PE, Phosphatidylethanolamine; PS, Phosphatidylserine; PG, Phosphatidylglycerol. Enzymes are represented by arrows, with their numbers in quadrants. Enzymes common nomenclature is used in the scheme. For an unambiguous identification, enzyme nomenclature (EC number system) is hereby given, along with further commonly used names: [1.1.1.94]- gpsA, glycerol-3-phosphate dehydrogenase; [1.1.5.3]- glpD, sn-glycerol-3-phosphate dehydrogenase; [2.3.1.15]- plsB, glycerol-3-phosphate O-acyltransferase; [2.3.1.274]- plsX, phosphate acyltransferase; [2.3.1.275]- plsY (ygiH), glycerol-3-phosphate acyltransferase; [2.3.1.51]- plsC (parF), 1-acyl-sn-glycerol-3-phosphate acyltransferase; [2.7.1.107]- dgkA, diacylglycerol kinase; [3.6.1.26]- cdh, CDP-diacylglycerol phosphotidylhydrolase, [2.7.7.41]- cdsA, (cds) CDP-diglyceride synthase; [EC:2.7.8.5]- pgsA, phosphatidylglycerophosphate synthetase; [3.1.3.27]- pgpA (yajN), phosphatidylglycerophosphatase A; [2.7.8.8]- pssA (pss), phosphatidylserine synthase; [3.1.1.32]- pldA, outer membrane phospholipase A; [4.1.1.65]- psd, phosphatidylserine decarboxylase; [3.1.1.4]- pldA, outer membrane phospholipase A; [2.3.1.40]- aas, 2-Acyl-sn-glycero-3-phosphoethanolamine O-acyltransferase; [3.1.1.5]- pldB, lysophospholipase L; [3.1.4.46]- glpQ (ugpQ), glycerophosphodiester phosphodiesterase. The T3SS-related shift in the phospholipid composition is mediated by *csrA* (B-D). The total abundance of identified phospholipids - phosphatidylglycerols (PGs; B) to phosphatidylethanolamines (PEs; C), lysophospholipids (LysoPLs; D) is presented. The expression of key enzymes responsible for the conversion of PLs to LysoPLs was evaluated by RT-qPCR: *pldA* (E) and *ygiH* (F). The data is presented as mean ± SE (N=4). **, p<0.01; ***, p<0.001.

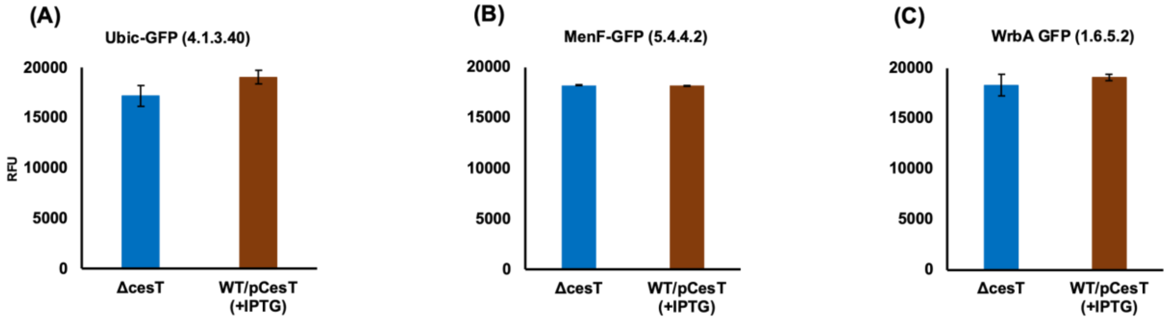


**Fig S5** No change noted in the expression of UbiC, MenF and WrbA following CesT accumulation. Quinone terpenoid biosynthesis genes were GFP labeled in EPEC null for *CesT* or overexpressing it. GFP fluorescence intensity was then measured. Data are presented as mean ± SE (N=3).

**TERPENOID QUINONE PATHWAY**

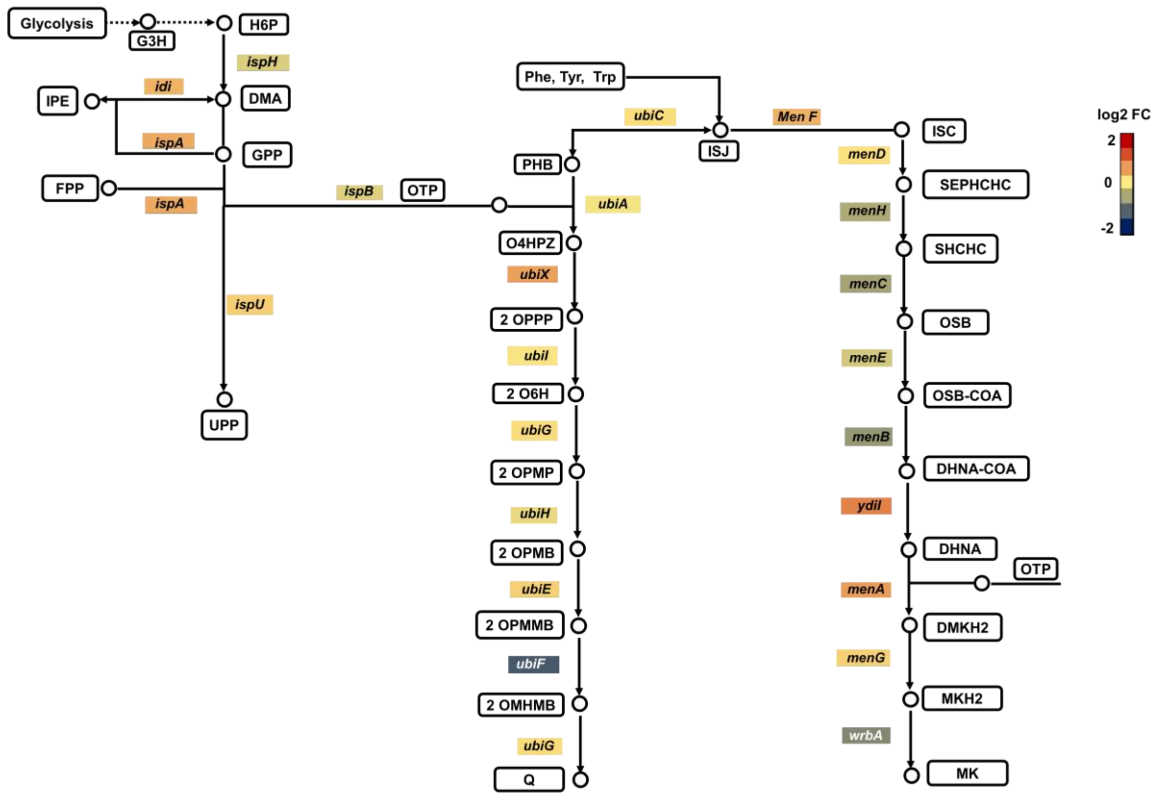


**Fig S6** Data mining in the Δ*csrA* transcriptome (1) suggests upregulation of undecaprenyl biosynthesis. Network was adapted from Kyoto Encyclopedia of Genes and Genomes (KEGG) for *Escherichia coli* O127:H6 E2348/69 (EPEC), and modified to include undecaprenyl, ubiquinone and menaquinone branches of terpenoid pathway. Lipids are represented by circles, with their names in colored quadrants. Enzyme are represented by arrows, with their names in colored quadrents. The color of quadrants represent the log2 fold change between the *csrA* mutant and the wild type (WT) strain. Lipids abbreviations: H6P, 1-Hydroxy-2-methyl-2-butenyl- 4-diphosphate; DMA, Dimethylallyl diphosphate; IPE, Isopentenyl diphosphate; GPP, Geranyl diphosphate; FPP, Farnesyl diphosphate; UPP, Undecaprenyl diphosphate; OTP, Octaprenyldiphosphate; PHB, Hydroxybenzoic acid; ISJ, Chorismate; O4HPZ, 4-Hydroxy-3-polyprenyl benzoate; 2OPPP, 2-Octaprenyl phenol; 2O6H, 2-Octaprenyl-6-hydroxyphenol; 2OPMP, 2-Octaprenyl-6-methoxy phenol; 2OPMB, 2-Octaprenyl-6-methoxy-1,4-benzoquinone; 2OPMMB, 2-Octaprenyl-3-methyl-6-methoxy-1,4-benzoquinone; 2OMHMB, 2-Octaprenyl-3-methyl-5-hydroxy-6-methoxy-1,4-benzoquinone; Q, Ubiquinone; ISC, Isochorismate; SEPHCHC, 2-Succinyl-5-enolpyruvyl-6-hydroxy-3-cyclohexene-1-carboxylate; SHCHC, 6-Hydroxy-2-succinylcyclohexa-2,4-diene-1-carboxylate; OSB, 2-Succinylbenzoate; OSB-COA, 2-Succinylbenzoyl-CoA; DHNA-CoA, 1,4-Dihydroxy-2-naphthoyl-CoA; DHNA, 1,4-Dihydroxy-2-naphthoate; DMKH2, Demethylmenaquinol; MKH2, Menaquinol; MK, Menaquinone. Enzymes common nomenclature is used in the scheme. For an unambiguous identification, enzyme nomenclature (EC number system) is hereby given, along with further commonly used names: [1.17.7.4]- ispH (yaaE, lytB), 4-hydroxy-3-methylbut-2-enyl diphosphate reductase; [2.5.1.1, 2.5.1.10]- ispA, farnesyl diphosphate synthase; [2.5.1.90]- ispB (cel, yhbD), all-trans-octaprenyl-diphosphate synthase; [2.5.1.31]- ispU (uppS, rth, yaeS), ditrans,polycis-undecaprenyl-diphosphate synthase; [4.1.3.40]-ubiC, chorismate lyase; [2.5.1.39] ubiA, 4-hydroxybenzoate octaprenyltransferase; [2.5.1.129]- ubiX (dedF), 3-octaprenyl-4-hydroxybenzoate carboxy-lyase; [1.14.13.240]- ubiI (visC), ), 2-octaprenyl phenol 6-hydroxylase; [2.1.1.222, 2.1.1.64]-ubiG (pufX, yfaB), bifunctional 3-demethylubiquinone-9 3-O-methyltransferase and 2-octaprenyl-6-hydroxyphenolmethylase; [1.14.13.-]- ubiH (acd, visB) 2-octaprenyl-6-methoxyphenol 4-hydroxylase; [2.1.1.201]- ubiE (yigO), bifunctional 2-octaprenyl-6-methoxy-1,4-benzoquinone methylase and demethylmenaquinone methyltransferase; [1.14.99.60]- ubiF (yleB), 2-octaprenyl-3-methyl-6-methoxy-1,4-benzoquinol oxygenase, 2-octaprenyl-3-methyl-6-methoxy-1,4-benzoquinol hydroxylase; [2.1.1.222, 2.1.1.64]- ubiG (pufX, yfaB) bifunctional 3-demethylubiquinone-9 3-O-methyltransferase and 2-octaprenyl-6-hydroxyphenol methylase; [5.4.4.2]- menF (yfbA), isochorismate synthase 2; [2.2.1.9]- menD, 2-succinyl-5-enolpyruvyl-6-hydroxy-3-cyclohexene-1-carboxylatesynthase; [4.2.99.20]- menH (yfbB), 2-

succinyl-6-hydroxy-2,4-cyclohexadiene-1-carboxylate synthase; [4.2.1.113]- menC, o-succinylbenzoate synthase; [6.2.1.26]- menE, o-succinylbenzoateCoA ligase; [4.1.3.36]- menB, 1,4-dihydroxy-2-naphthoyl-CoA synthase; []- ydiI (menI), 1,4-dihydroxy-2-naphthoyl-CoA hydrolase; [2.5.1.74]- menA (yiiW), 1,4-dihydroxy-2-naphthoate octaprenyltransferase; [2.1.1.163, 2.1.1.201]- ubiE (menG, yigO), demethylmenaquinone methyltransferase /2-methoxy-6-octaprenyl-1,4-benzoquinol methylase; [1.6.5.2]- wrbA(ytfG), quinoneoxidoreductase.

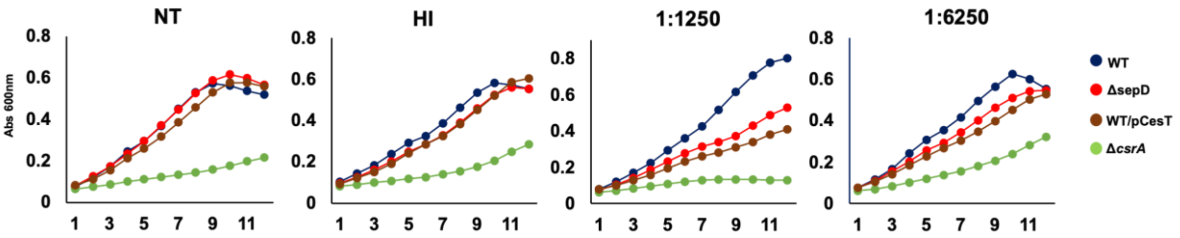


**Fig S7** Following T3SS activation EPEC exhibit lower resistance to serum bactericidal components. EPEC strains: wild type, Δ*sepD*, Δ*csrA* and WT/pCesT were grown overnight in media containing human sera of different dilutions, or no serum (non-treated: NT), or heat inactivated sera (HI) at 37$^0$ C in a Spark 10M microplate reader. Bacterial growth kinetics was monitored at hourly intervals for 12 hours. The data represented are mean value of N=3 replicates.

.

**Table S1** Identified lipids taken from the lipidomics analysis

| ID | Adduct | Empirical Formula | FC (ΔsepD/WT) |
|---|---|---|---|
| PG (14:0 /17:0) | 731.47(M+Na) | $C_{37}H_{73}O_{10}P$ | 0.52 |
| PG (12:0 /15:0) | 675.42 (M+Na) | $C_{33}H_{65}O_{10}P$ | 0.53 |
| PG (14:0 OH/19:0) | 735.508 (M+H-$H_2O$) | $C_{39}H_{77}O_{11}P$ | 0.37 |
| PG (14:0 OH/15:0) | 712.47 (M+$NH_4$) | $C_{35}H_{69}O_{11}P$ | 0.52 |
| PG (14:0 /14:0) | 667.45 (M+H) | $C_{34}H_{67}O_{10}P$ | 0.51 |
| PG (16:0/16:0) | 740.54 (M+$NH_4$) | $C_{38}H_{75}O_{10}P$ | 0.57 |
| PG (16:0/17:0) | 757.49 (M+Na) | $C_{39}H_{75}O_{10}P$ | 0.46 |
| PG(17:0/17:0) | 747.50 (M+H) | $C_{40}H_{75}O_{10}P$ | 0.51 |
| PG(17:1/17:2) | 767.48 (M+Na) | $C_{40}H_{73}O_{10}P$ | 0.63 |
| PG (14:0/16:0) | 717.46 (M+Na) | $C_{33}H_{71}O_{10}P$ | 0.54 |
| PG (14:0 OH/18:1) | 737.48 (M+H) | $C_{38}H_{73}O_{11}P$ | 0.38 |
| PG (12:0/12:0) | 633.38 (M+ Na) | $C_{30}H_{59}O_{10}P$ | 1.02 |
| PGP (14:0 /16:1) | 1547.91 (2M+H) | $C_{36}H_{70}O_{13}P_2$ | 0.43 |
| PS(18:0) | 508.30 (M+H-$H_2O$) | $C_{24}H_{48}NO_9P$ | 2.88 |
| PG (16:1) | 465.25(M+H-$H_2O$) | $C_{22}H_{43}O_9P$ | 2.53 |
| PE(16:1) | 452.27 (M+H) | $C_{21}H_{42}NO_7P$ | 2.53 |
| PE(18:1) | 480.30 (M+H) | $C_{23}H_{46}NO_7P$ | 0.83 |
| PE (16:1) | 452.28(M+H) | $C_{21}H_{42}O_7P$ | 2.68 |
| PS(19:0) | 522.31 (M+H-$H_2O$) | $C_{25}H_{50}NO_9P$ | 2.20 |
| PE(19:0) | 494.32 (M+H) | $C_{24}H_{48}NO_7P$ | 3.06 |
| PE(17:1) | 488.27 (M+Na) | $C_{22}H_{44}NO_7P$ | 3.13 |
| PE(16:0) | 454.29 (M+H) | $C_{21}H_{44}NO_7P$ | 1.40 |
| PE(19:0) | 518.32 (M+Na) | $C_{24}H_{48}NO_7P$ | 1.07 |
| PG(14:1) | 477.22 (M+Na) | $C_{20}H_{39}O_9P$ | 14.17 |
| PA(18:1) | 419.26 (M+H) | $C_{21}H_{41}O_7P$ | 2.63 |
| PE(17:0) | 466.29 (M+H) | $C_{22}H_{44}NO_7P$ | 1.13 |
| PG(12:0) | 451.20(M+Na) | $C_{18}H_{37}O_9P$ | 1.56 |
| PG(15:0) | 493.25 (M+Na) | $C_{21}H_{43}NO_9P$ | 21.58 |
| PS(18:1) | 506.28 (M+H-$H_2O$) | $C_{24}H_{46}NO_9P$ | 92.59 |
| PG(18:1) | 533.28 (M+Na) | $C_{24}H_{47}O_7P$ | 1.95 |
| PA (18:2) | 457.23(M+Na) | $C_{21}H_{41}O_7P$ | 5.12 |
| PE(17:1) | 466.29 (M+H) | $C_{22}H_{44}NO_7P$ | 1.92 |
| PG(12:0) | 429.21 (M+H) | $C_{18}H_{37}O_9P$ | 4.46 |
| PG (15:0) | 491.23 (M+Na) | $C_{21}H_{41}O_9P$ | 94.26 |
| PG (13:0) | 407.21 (M+H-2$H_2O$) | $C_{19}H_{39}O_9P$ | 0.54 |
| PS(17:1) | 510.28 (M+H) | $C_{23}H_{44}NO_9P$ | 90.46 |
| PS(17:0) | 510.28 (M+H) | $C_{23}H_{44}NO_9P$ | 0.67 |
| PA(17:1) | 423.24 (M+H) | $C_{20}H_{39}O_7P$ | 1.21 |
| PG(16:0) | 507.27 (M+Na) | $C_{22}H_{45}O_9P$ | 164677.08 |
| PG(16:0) | 467.27 (M+H-$H_2O$) | $C_{22}H_{45}NO_9P$ | 0.08 |
| PE(17:0) | 512.27(M+2Na-H) | $C_{22}H_{46}NO_7P$ | 26.67 |
| PG(14:0) | 457.25 (M+H) | $C_{20}H_{41}O_9P$ | 5.50 |
| PE(18:0) | 504.31 (M+Na) | $C_{23}H_{48}NO_7P$ | 0.18 |
| PE(12:0) | 420.21 (M+Na) | $C_{17}H_{36}NO_7P$ | 1.31 |
| PS(14:0) | 470.25 (M+H) | $C_{20}H_{40}NO_9P$ | 0.00 |
| PG(18:1) | 510.3 (M+$NH_4$) | $C_{24}H_{47}O_9P$ | 0.42 |
| PG (15:1) | 469.26 (M+H) | $C_{21}H_{41}O_9P$ | 0.00 |
| CL(64:1) | 1333.94 (M+H-$H_2O$) | $C_{73}H_{140}O_{17}P_2$ | 0.57 |
| CL(69:0) | 1418 (M+H) | $C_{78}H_{146}O_{17}P_2$ | 1.72 |
| CL(68:1) | 1402.01(M+H-$H_2O$) | $C_{81}H_{150}O_{17}P_2$ | 2.27 |
| CL(70:1) | 1453.07 (M+$NH_4$) | $C_{79}H_{152}O_{17}P_2$ | 0.63 |
| CL(68:4) | 1434.00 (M+$CH_3OH$+H) | $C_{77}H_{142}O_{17}P_2$ | 1.94 |
| CL(68:2) | 1464.98(M+Acn+Na) | $C_{77}H_{142}O_{17}P_2$ | 4.39 |
| CL(70:2) | 1416.01 (M+H-$H_2O$) | $C_{79}H_{150}O_{17}P_2$ | 2.16 |
| CL(63:3) | 1417.91 (M+2Na-H) | $C_{75}H_{138}O_{17}P_2$ | 2.36 |
| CL(66:2) | 1440.98 (M+Acn+Na) | $C_{75}H_{142}O_{17}P_2$ | 1.76 |
| CL(72:2) | 1440.03 (M+H-$H_2O$) | $C_{81}H_{156}O_{17}P_2$ | 1.20 |
| PE(17:0/14:0) | 676.49 (M+H) | $C_{36}H_{70}NO_8P$ | 0.88 |

| | | | |
|---|---|---|---|
| PE (17:0/18:1) | 730.53 (M+H) | $C_{39}H_{76}NO_8P$ | 0.27 |
| PE (17:0/15:0) | 690.50 (M+H) | $C_{39}H_{74}NO_8P$ | 1.80 |
| PE (16:1/18:1) | 716.51 (M+H) | $C_{39}H_{74}NO_8P$ | 3.07 |
| PE (19:0/10 OH) | 664.45 (M+H) | $C_{34}H_{66}NO9P$ | 2.57 |
| PE (16:0/12:0) | 636.46 (M+H) | $C_{33}H_{66}NO_8P$ | 2.96 |
| PE (19:0/14:0) | 726.50( M+Na) | $C_{41}H_{78}NO_8P$ | 2.40 |
| PE (12:0/16:0) | 636.46(M+H) | $C_{36}H_{68}NO_8P$ | 5.60 |
| PE (16:0/19:0) | 732.55 (M+H) | $C_{40}H_{78}NO_8P$ | 0.52 |
| PE (17:0/10:0) | 620.42 (M+H) | $C_{32}H_{62}NO_8P$ | 1.68 |
| PE (17:0/17:0) | 716.51 (M+H) | $C_{40}H_{76}NO_8P$ | 1.68 |
| PE (19:0/16:1) | 730.53 (M+H) | $C_{39}H_{74}NO_8P$ | 0.85 |
| PE(17:0/16:1) | 719.53 (M+NH<sub>4</sub>) | $C_{38}H_{72}NO_8P$ | 0.43 |
| PE(18:1/14:0 OH) | 680.48 (M+H) | $C_{35}H_{70}NO_9P$ | 0.57 |
| PE(19:0/17:0) | 744.54 (M+H) | $C_{41}H_{78}NO_8P$ | 1.67 |
| PE(19:0/18:1) | 758.56 (M+H) | $C_{42}H_{80}NO_8P$ | 0.76 |
| PE(14:0/17:0) | 678.50 (M+H) | $C_{36}H_{72}NO_8P$ | 2.89 |
| PE(18:1/19:0) | 782.5618 (M+Na) | $C_{42}H_{82}NO_8P$ | 1.57 |
| PE (16:1/19:0) | 730.54 (M+H) | $C_{40}H_{78}NO_8P$ | 0.60 |
| PE(19:0/16:0) | 732.54 (M+H) | $C_{40}H_{78}NO_8P$ | 2.39 |
| PE(16:0/18:1) | 718.53 (M+H) | $C_{39}H_{76}NO_8P$ | 1.26 |
| PE (18:1/18:1) | 744.55 (M+H) | $C_{41}H_{78}NO_8P$ | 1.18 |
| PE(16:1/14:1) | 660.45 (M+H) | $C_{35}H_{66}NO_8P$ | 0.83 |
| PE (15:0/15:0) | 642.44 (M+H-$H_2O$) | $C_{35}H_{66}NO_8P$ | 1.94 |
| PE (16:1/12:0) | 656.42 (M+Na) | $C_{33}H_{64}NO_8P$ | 0.66 |
| PE (19:0/14:0 OH) | 720.51 (M+H) | $C_{38}H_{74}NO_9P$ | 1.40 |
| PE(16:0/17:0) | 704.51 (M+H) | $C_{38}H_{74}NO_8P$ | 0.88 |
| PE (16:0/14:0 OH) | 718.43 (M+K) | $C_{35}H_{70}NO_9P$ | 0.12 |
| PE(19:1/15:0) | 735.56 (M+$NH_4$) | $C_{39}H_{76}NO_8P$ | 0.79 |
| PE(17:0/14:0) | 676.48 (M+H) | $C_{38}H_{74}NO_8P$ | 0.64 |
| PE(10:0/18:1) | 634.43 (M+H) | $C_{38}H_{76}NO_9P$ | 0.01 |
| PE(12:0(OH)/16:1) | 650.43 (M+H) | $C_{35}H_{70}NO_9P$ | 0.78 |
| PE(19:1/18:1) | 775.59(M+$NH_4$) | $C_{4}OH_{68}O_7P_2$ | 0.78 |
| PE(36:0) | 748.57 (M+H) | $C_{41}H_{82}NO_8P$ | 3.00 |
| PE(35:0) | 776.51 (M+2Na-H) | $C_{40}H_{78}NO_8P$ | 0.76 |
| PE(29:1) | 692.42 (M+2Na-H) | $C_{34}H_{66}NO_8P$ | 1.66 |
| PE(36:0) | 792.56 (M+2Na-H) | $C_{41}H_{82}NO_8P$ | 2.88 |
| PE(19:1/18:0) | 760.57 (M+H) | $C_{42}H_{82}NO_8P$ | 2.38 |
| PE(37:1) | 780.56 (M+Na) | $C_{42}H_{80}NO_8P$ | 1.94 |
| PE(36:0) | 792.54 (M+2Na-H) | $C_{41}H_{82}NO_8P$ | 1.68 |
| PE(38:0) | 820.58(M+2Na-H) | $C_{43}H_{86}NO_8P$ | 2.57 |
| Octaprenyl diphosphate | 723.44 (M+H) | $C_{42}H_{82}NO_8P_2$ | 6.28 |
| Glycosyl undecaprenyl phosphate | 1040.74 (M+$CH_3OH$) | $C_{61}H_{100}O_9P$ | 12.07 |
| Glycosyl undecaprenyl phosphate | 1072.74 (M+Acn+Na) | $C_{61}H_{100}O_9P$ | 1.57 |
| Undecaprenyl diphosphate | 909.63 (M+H-$H_2O$) | $C_{55}H_{91}O_7P_2$ | 2.01 |
| Undecaprenyl phosphate | 910.67 (M+Acn+Na) | $C_{55}H_{89}O_7P$ | 1.19 |
| Undecaprenyl diphsophate | 927.63 (M+H) | $C_{55}H_{92}O_7P_2$ | 0.84 |
| Glycosyl undecaprenyl phosphate | 1028.70 (M+Na) | $C_{61}H_{99}NO_8P$ | 64.65 |
| Menaquinone 8 | 739.54 (M+Na) | $C_{51}H_{72}O_2$ | 0.82 |
| Menaquinone 8 | 1456.09 (2M+Na) | $C_{51}H_{72}O_2$ | 0.99 |
| Ubiquinone 6* | 591.44 (M+H) | $C_{39}H_{58}O_4$ | 0.82 |
| Ubiquinone 7* | 659.50 (M+H) | $C_{44}H_{66}O4$ | 0.59 |
| Ubiquinone 8 | 727.56 (M+H) | $C_{49}H_{74}O_4$ | 0.52 |
| Octaprenyl phenol | 656.58 (M+$NH_4$) | $C_{46}H_{70}O$ | 6.48 |
| Methoxy octaprenyl benzoquinol | 1433.09 (2M+Acn+Na) | $C_{47}H_{72}O_3$ | 0.32 |

PG, Phosphatidylglycerol; PE, phosphatidylethanolamine; PS, phosphatidylserine; PA, phosphatidic acid; CL, cardiolipin.The lipid species were identified by $MS^E$. The Data shown in the table represent the fold change calculated from the abundance of each lipid in the Δ*sepD* compared to the wild type strain.
*, an intermeiate ubiquinone

**Table S2** Strains and plasmids used for cloning

| Name (Number in our collection) | Description | Reference |
|---|---|---|
| **E2348/69 (1)** | EPEC wild type isolate O127:H6 | (2) |
| **EM2018** | E2348/69 Δ*cesT*::kn | (2) |
| **NN6237** | Dh5a containing pNN6237 (pCesT) | (3) |
| **NN5898** | E2348/69 Δ*csrA*::cm | (3) |
| **5768** | XTL634 as template for tet-sacB cassette | (4) |
| **A8695** | NN6237 containing wrbA-gfp translational fusion | |
| **A8696** | NN6237 containing idi-gfp translational fusion | |
| **A8698** | NN6237 containing ispU-gfp translational fusion | |
| **A8700** | NN6237 containing menF-gfp translation fusion | |
| **A8704** | NN6237 containing ubiC-gfp translational fusion | |
| **pKD46 (p811)** | Contains the λ red genes. | (5) |
| **pCesT (pNS6237)** | A pSA10 derivative containing cesT between EcoRI and SalI sites, cm resistance | (3) |
| **pCesT (pSK6194)** | A pSA10 derivative containing cesT between EcoRI and SalI sites, amp resistance | (3) |
| **EM4620** | EPEC nleA-gfp | (2) |

**Table S3** Primer sequence for GFP fusions in EPEC

| Gene fusions | Primer Nos. in collection | Primer Names | Sequences (5'-3') |
|---|---|---|---|
| **idi-gfp** | 4317 | Idi_GFP_F | GCGACAAATCGCGAAGCCAGAAAACGATTATCTGCATTTACCCAGCTTAAAG GTGGTACCCGTAAAGGAGAAGAAC |
| | 4318 | idi_GFP_R | GTAAAATAAGCATTACGTTATGCTCACAACCCCGGCAACCGTCGGGGTTTTT ATTATTTGTATAGTTCATCCATGCC |
| **idi-tet-sacB** | 4301 | idi_Tet_F | GCGACAAATCGCGAAGCCAGAAAACGATTATCTGCATTTACCCAGCTTAAATA ATCCTAATTTTTGTTGACACTCTATC |
| | 4302 | idi_SacB_R | GTAAAATAAGCATTACGTTATGCTCACAACCCCGGCAACCGTCGGGGTTTTAT CAAAGGGAAAACTGTCCAT |
| **menF-gfp** | 4327 | menF_GFP_F | GGCAGGAAATCGACAACAAAGCGGCAGGGCTGCGTACTTTATTACAAATGGA AGGTGGTACCCGTAAAGGAGAAGAAC |
| | 4328 | menF_GFP_R | GGAGCTACAAAAATAGATTATTGATATGAATCGGTAATGATGCGACTCATTAC TATTATTTGTATAGTTCATCCATGCC |
| **menF-tet-sacB** | 4307 | menF_Tet_F | GCAGGAAATCGACAACAAAGCGGCAGGGCTGCGTACTTTATTACAAATGGAA TAGTCCTAATTTTTGTTGACACTCTATC |
| | 4308 | menF_SacB_R | GGAGCTACAAAAATAGATTATTGATATGAATCGGTAATGATGCGACTCATTAA TCAAAGGGAAAACTGTCCATA |
| **ubiC-gfp** | 4311 | ubic_GFP_F | AAGCGGTAAACCGCTGTTGCTAACAGAACTGTTTTTACCGGCGTCACCGTTGT ACGGTGGTACCCGTAAAGGAGAAGAAC |
| | 4312 | ubic_GFP_R | GAAACGCCAGCAGCTTATTCTGCGTCAGACTCCACTCCATATTTTTTTCCTCTT ATTATTTGTATAGTTCATCCATGCC |
| **ubiC-tet-sacB** | 4293 | ubic_Tet_F | AAGCGGTAAACCGCTGTTGCTAACAGAACTGTTTTTACCGGCGTCACCGTTGT ACTAATCCTAATTTTTGTTGACACTCTATC |
| | 4294 | ubic_SacB_R | GAAACGCCAGCAGCTTATTCTGCGTCAGACTCCACTCCATATTTTTTTCCTCAT CAAAGGGAAAACTGTCCATA |
| **ispU-gfp** | 4313 | IspU_GFP_F | CTTTGCTAATCGAGAGCGTCGTTTCGGCGGCACCGAGCCCGGTGATGAAACAG CCGGTGGTACCCGTAAAGGAGAAGAAC |
| | 4314 | IspU_GFP_R | GGTATTAACACAAAAGCAGATATCAGGCGATACTTCAGCAAAAGCGACCCCC ATCATTATTTGTATAGTTCATCCATGCC |
| **ispU-tet-sacB** | 4291 | IspU_Tet_F | GCTAATCGAGAGCGTCGTTTCGGCGGCACCGAGCCCGGTGATGAAACAGCCT GATCCTAATTTTTGTTGACACTCTATC |
| | 4292 | IspU_Sac_B | GGTATTAACACAAAAGCAGATATCAGGCGATACTTCAGCAAAAGCGACCCCC AATCAAAGGGAAAACTGTCCATA |
| **wrbA-gfp** | 4323 | wrbA_GFP_F | GCTCGTTATCAAGGGGAATATGTCGCAGGTCTGGCAGTTAAACTTAACGGCGG TGGTACCCGTAAAGGAGAAGAAC |
| | 4324 | wrbA_GFP_R | GACGTGATGAGCTTTCGCTTCTTGAGTTGGCATGCGTATCCTCCTGTTGAAGAT TATTATTTGTATAGTTCATCCATGCC |
| **wrbA-tet-sacB** | 4299 | wrbA_Tet_F | GCTCGTTATCAAGGGGAATATGTCGCAGGTCTGGCAGTTAAACTTAACGGCTA ATCCTAATTTTTGTTGACACTCTATC |
| | 4300 | wrbA_SacB_R | GACGTGATGAGCTTTCGCTTCTTGAGTTGGCATGCGTATCCTCCTGTTGAAGAA TCAAAGGGAAAACTGTCCATA |

**Table S4** Primers used for PCR confirmation of GFP fusions

| Genes | Primer Nos. in collection | Primer Names | Sequences (5'-3') |
|---|---|---|---|
| **ispU** | 4482 | ispUcnfm-F | CGTTGGCTGTTTAACGCCTG |
| | 4481 | ispU-R2 | CTGTCCTTAATTCCTGCTTC |
| **menF** | 4479 | menFcnfm-F | GAAGCAGGCTGTTTATCGTG |
| | 4478 | menF-R2 | CCACTATTTGTGCCTGTTGC |
| **ubiC** | 4473 | ubiCcnfm-F | TGTTCTGGTCTTTACCACGG |
| | 4472 | ubiC-R2 | CCAGTAACTCATTGCCAGCC |
| **wrbA** | 4476 | wrbAcnfm-F | GCATGCGTCATTGATTAGCC |
| | 4475 | wrbA-R2 | GATGTTCATCAGTTACAGCG |
| **idi** | 4447 | idi-cnfmF | AATCCTGTTTGGCATGGTGC |
| | 4360 | idi-R | ACCAACAGCCATACCATCCG |

**Table S5** Primers used for QPCR

| Genes | Primer Names | Sequences (5'-3') |
|---|---|---|
| **pldA** | pldA-F1 | GGTTGTTGCCGGTGTTTATG |
| | pldA-R1 | CCAGGATTTCTGTGTGTAGGAG |
| **ygiH** | ygiH-F1 | CTCCATTTCCAGTGCCATTCT |
| | ygiH-R1 | GCGACGAGTAACCACTCAATAG |
| **Aas** | aas-F1 | GCCAACCCGTATGACTTCTATC |
| | aas-R1 | CATTTCCAGCGACACCATTTC |
| **rrsB** | rrsB-F1 | CAG AGATGAGAATGTGCCTTCGGG |
| | rrsB-R1 | CCGCTGGCAACAAAGGATAAG G |

**Table S6** The linear gradient for the lipidomics analyses

| Time | Flow (mL/min) | %A Water (0.1 FA) | %B Acetonitrile (0.1 FA) | %C (Isopropanol) |
|---|---|---|---|---|
| **0.0-1.0** | 0.4 | 60 | 40 | 0 |
| **1.0-5.0** | 0.4 | 30 | 70 | 0 |
| **5.0-8.0** | 0.4 | 24 | 40 | 36 |
| **8.0-9.0** | 0.4 | 20 | 35 | 45 |
| **9.0-12.0** | 0.4 | 18.4 | 33 | 48.6 |
| **12.0-17** | 0.4 | 12 | 25 | 63 |
| **17.0-25.50** | 0.4 | 0.4 | 10.5 | 89.1 |
| **26.0-35.0** | 0.4 | 60 | 40 | 0 |

### References


.1 Berndt V, Beckstette M, Volk M, Dersch P, Bronstrup M. Metabolome and transcriptome-wide effects of the carbon storage regulator A in enteropathogenic Escherichia coli. Sci Rep. 2019;9(1):138.

.2 Katsowich N, Elbaz N, Pal RR, Mills E, Kobi S, Kahan T ,et al. Host cell attachment elicits posttranscriptional regulation in infecting enteropathogenic bacteria. Science. 2017;355(6326):735-9.

.3 Jeucken A, Molenaar MR, van de Lest CHA, Jansen JWA, Helms JB, Brouwers JF. A Comprehensive Functional Characterization of Escherichia coli Lipid Genes. Cell Rep. 2019;27(5):1597-606 e2.

.4 Li XT, Thomason LC, Sawitzke JA, Costantino N, Court DL. Positive and negative selection using the tetA-sacB cassette: recombineering and P1 transduction in Escherichia coli. Nucleic Acids Res. 2013;41(22):e204.

.5 Datsenko KA, Wanner BL. One-step inactivation of chromosomal genes in Escherichia coli K-12 using PCR products. Proc Natl Acad Sci U S A. 2000;97(12):6640-5.
